# Circadian phosphotimer generates time cues through PER dimerization-mediated trans-phosphorylation

**DOI:** 10.1101/2025.10.01.679838

**Authors:** Kwangjun Lee, Jiyoung Park, Junghyun Lee, Jun Ho, Christian Hong, Sookkyung Lim, Choogon Lee

## Abstract

The circadian clock generates autonomous molecular rhythms that regulate sleep cycles, chronotypes, daily physiology, and expression patterns of almost the entire genome. Temporal phosphorylation of PERIOD (PER) proteins creates a phosphotimer that is critical for clock function. Although previous studies show that PER stably binds the casein kinase CK1, cis-acting phosphorylation of PER by CK1 is not compatible with slow PER phosphorylation extended over 24 hrs. So how can PER phosphorylation be programmed in a slow and controlled manner? Here, we show that the timing cues are encoded by PER-PER dimerization-mediated trans-phosphorylation, which enables the necessary time delay in phosphorylation and slow phosphorylation. When PER dimerization is disrupted, PER phosphorylation and circadian rhythms are severely compromised. In mouse models with point mutations in the PER dimerization domain, circadian period is shortened to ∼20 hrs, and the phase of wake/sleep cycles is dramatically advanced, switching a nocturnal animal to a half diurnal animal.

## Introduction

The circadian clock is the master regulator of circadian rhythms including sleep/wake cycles ^1,2^. General physiology, behavior, and health are profoundly impacted by circadian rhythms—and by their disruption ^3–6^. Decades of prior work have revealed that the clock is built on a core negative feedback loop that is cell autonomous, involving transcriptional and post-translational regulation of several essential clock genes ^6,7^. In the feedback loop, CLOCK:BMAL1 (the activator complex) drives transcription of two redundant pacemaker *Period* (*Per*) genes, *Per1* and *Per2*, and two partially redundant *Cryptochrome* (*Cry*) genes, *Cry1* and *Cry2*, along with many other clock-controlled genes (ccgs). PER acts as a scaffolding protein to form an inhibitory complex that includes CRY as well as casein kinases CK1δ and/or CK1ε— functionally redundant kinases collectively referred to as CK1. This inhibitory complex suppresses the activity of CLOCK:BMAL1, thereby completing the feedback loop ^8–11^. Both mathematical modeling and experimental evidence have shown that a proper stoichiometric relationship among clock components is critical for a robust feedback loop, with *Per* acting as the rate-limiting component in the system ^12–15^.

This feedback loop also depends on a phosphotimer to mediate timely nuclear translocation of the inhibitory complex and the interaction between the activator and the inhibitory complex ^13,16–19^. Circadian parameters such as period and phase (e.g., activity or sleep onset) are ultimately determined by PER oscillations in nuclear entry/accumulation and degradation—processes temporally controlled by CK1-mediated phosphorylation ^13,18,20–22^. This phosphotimer mechanism has been supported by numerous in vitro and genetic studies, some of which linked specific single nucleotide polymorphisms (SNPs) to altered circadian rhythms ^21,23–25^. The best studied is a SNP in the human *Per2* gene that disrupts a phosphorylation site (S662G) and causes familial advanced sleep-phase (FASP) syndrome ^23,26^. This SNP alters PER phosphorylation kinetics, and the resulting altered rhythms can be recapitulated in cultured cells and mice with the same mutation ^21,26–28^.

PER proteins have several key domains and motifs, including the PAS dimerization domain, the CK1-binding domain (CKBD), the CRY-binding domain (CBD), a degron motif, and a nuclear localization signal (NLS) ^19,27,29,30^. The FASP mutation resides within the central region of the CKBD. Through these domains and motifs, PER recruits several key clock components, which then regulate PER phosphorylation, subcellular location, and activity. These properties, in turn, modulate the fate of other components in the PER complex. It has been shown that stable interaction between PER and CK1 through the CKBD is critical for PER phosphorylation ^19,31,32^, which intuitively suggested cis-acting phosphorylation in the PER:CK1 complex. This assumption of cis-phosphorylation could not explain readily why there is a long delay in phosphorylation in early phase of PER accumulation and slow phosphorylation kinetics ^17^. Amino acid (AA) indels at the FASP locus (middle of CKBD) dramatically affect period by modulating phosphorylation speed, despite overall phosphorylation levels remaining intact ^21,33^. However, a recent study suggested that PER interaction with CK1 through CKBD may not be critical for the circadian clock ^32^. Specifically, disruption of the C-terminal side of the PER2 CKBD through mutation of two AAs severely impairs PER2 phosphorylation, yet only marginally alters behavioral rhythms, even in a *Per1* KO background. Since the central dogma of the circadian clock holds that oscillations of at least some key components are necessary to generate a circadian negative feedback loop, these findings call for reconciliation with the current understanding of the dogma.

The PAS domain mediates homo- or hetero-dimerization of PER proteins with one another ^17,34^. While the necessity of other domains such as CKBD and CBD is intuitive—they are needed for CK1 and CRY complex formation with PER—the significance of PAS in generating time cues remains unclear. In our previous studies ^35,36^, we suggested that PER dimerization or multimerization should be an essential feature in the circadian feedback loop because it enables individual pacemaker protein molecules to behave collectively through PER-PER cooperativity. The core mechanism of the circadian clock is the temporal and robust repression and de-repression of the activator complex by the inhibitor complex, which is driven by temporal trafficking of the PER-containing complex from the cytosol to the nucleus, followed by its degradation. Given that *Per* is transcribed and translated over ∼10 hrs in the cytosol, this creates great spatiotemporal heterogeneity among thousands of PER molecules within a single clock cell ^37^. To compensate for this variability, collective behavior among PER molecules is likely necessary. PAS can dimerize through two interfaces, PAS-A and PAS-B, which are the only known PER-PER dimerization interfaces, suggesting that PAS may play a crucial role in coordinating this collective behavior.

In the present study, we demonstrate that circadian time cues are generated in a nonlinear manner by the kinetics of PER dimerization-induced trans-phosphorylation, with the PAS domain responsible for dimerization. The PAS-mediated phosphorylation dynamics create an initial delay and drive precise progression of the circadian time-keeping phosphotimer. The kinetics of trans-phosphorylation are dictated by the efficiency of PAS:PAS dimerization; thus, any disruption in the PAS dimerization interface results in a dramatic alteration in the phase and the period of circadian rhythms. Our findings provide direct insights into the diverse pathogenesis and chronotypes that may arise from human SNPs in the PAS domain, as modeled in mice with similar SNPs.

## Results

### PAS domain in PER is critical for PER phosphorylation and circadian rhythms

As alluded to above, we hypothesized that multiple domains in PER may contribute to PER phosphorylation in a cooperative manner. To identify critical domains for the phosphotimer function in PER, we disrupted all major domains in PER by creating random AA indels using CRISPR in U2OS reporter cells, a human cell line with a luciferase (Luc) reporter widely used as a circadian clock model. We then assessed alterations in circadian bioluminescence rhythms and PER phosphorylation, aiming to elucidate the mechanistic relationships underlying changes in temporal signaling (Fig S1A) ^33^. Based on our prior prediction, we first evaluated the function of the PAS domain in relation to phosphorylation and circadian rhythms (Fig 1A). We focused initially on PER1 PAS, as the preliminary screening process did not require molecular analysis. Dramatic phase shifts in the bioluminescence rhythms can be induced when *Per1* gene is compromised ^33^, allowing candidate clones to be selected based solely on altered phase in the bioluminescence rhythms (Fig 1B). From these clones, mutant clones with AA indels (in-frame mutations) in PER1 PAS-A were selected by immunoblotting (Fig 1C). One such mutant clone (#2-12) was isolated in U2OS-*Bmal1-Luc* cells. This clone exhibited a severely hypophosphorylated PER1 and was found to lack nine AAs in the PAS-A domain (Fig 1D), a motif that serves as a linker connecting the dimerization interface in PAS-A to the N-terminus (Fig 1E and F). The absence of this linker is therefore expected to disrupt PER1 dimerization. Two additional AA indel clones in PAS-A were subsequently isolated in U2OS-*Bmal1-Luc*/*Per2 KO* cells, and both mutant PER1 proteins were also severely hypophosphorylated (Fig S1B). Deletion of the *Per2* gene in the first clone (#2-12) resulted in arrhythmicity, indicating that the mutant PER1 is nonfunctional in the absence of PER2 (Fig S1C). The mutant PER1 was constitutively hypophosphorylated over a circadian cycle, while PER2 phosphorylation remained intact in the same clone (Fig 1G and H). The mutant PER1 was never hyperphosphorylated, from its de novo synthesis to degradation (Fig S1D and E). Previous studies have shown that PER phosphorylation is regulated by a dynamic balance between CK1 and protein phosphatase 1 (PP1) ^18^. Thus, the mutant PER1 may appear hypophosphorylated because it becomes a more favorable substrate for PP1 relative to CK1, resulting in a persistently hypophosphorylated steady state. To test this idea, both control and mutant cells were treated with Calyculin A (CA) to inhibit PP1. As previously shown, wt PER1 became excessively hyperphosphorylated beyond its normal level of hyperphosphorylation by the treatment, whereas the mutant PER1 remained severely hypophosphorylated. This suggests that phosphorylation is inherently defective in the mutant PER1 (Fig 1I). These findings are counterintuitive because the CKBD is far downstream from the deletion motif, meaning that the small deletion should not directly affect binding with CK1. Indeed, CK1 binding to the mutant PER1 was not significantly altered compared to wt PER1 (Fig 1J). When wt and the PAS mutant *Per1* were transfected into cells, wt PER1 became hyperphosphorylated, whereas the mutant PER1 was hypophosphorylated (Fig 1K). When they were co-expressed with wt PER1-Luc and immunoprecipitated with anti-Luc antibody, only wt PER1 was efficiently co-purified (Fig 1K). Consistent with its phosphorylated status, the mutant PER1 was predominantly localized in the cytoplasm (Fig S1F). The CLOCK protein is typically hyperphosphorylated by nuclear PER:CK1 complex; however, CLOCK was hypophosphorylated in the mutant clone lacking *Per2*, aligning with the cytoplasmic localization of the mutant PER1 (Fig S1G).

**Fig. 1.**
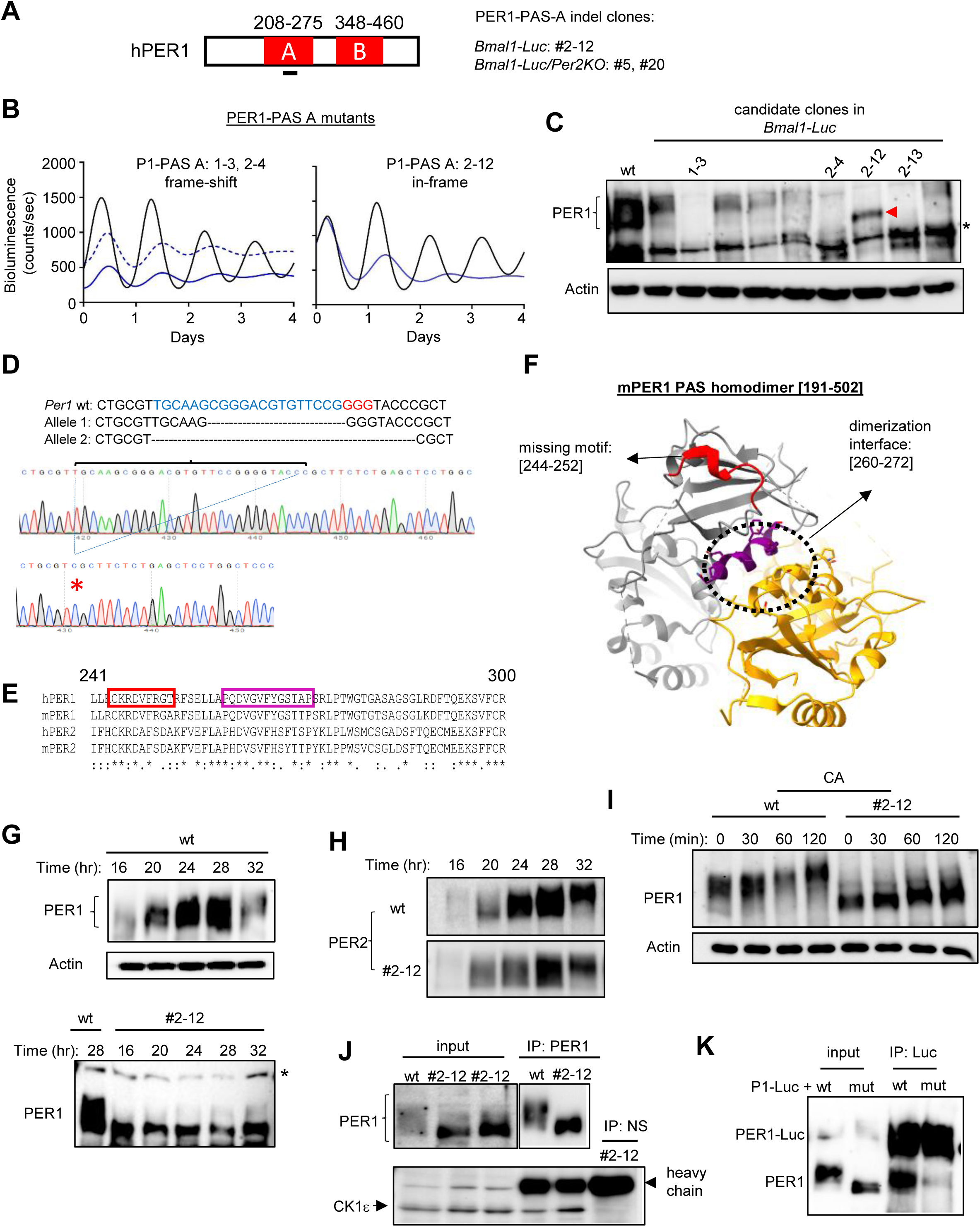
PAS-A domain in PER1 is critical for PER1 phosphorylation and circadian rhythms. (A) There are two PAS domains in PER, PAS-A and PAS-B. AA indel clones in PER1-PAS-A were selected in two different transgenic reporter U2OS cell lines: *Bmal1-Luc*, in which the *Bmal1* promoter drives luciferase expression, and *Bmal1-Luc/Per2KO*, in which the same reporter is present together with a *Per2* knockout genotype. (B, C) As we have done previously, we transfected cells with our all-in-one CRISPR plasmid and selected and sorted to single cells by FACS ^33^. Candidate clones were selected based on altered circadian phase (B) before they were subjected to immunoblotting for measuring phosphorylation. N=3 each in (B). (C). Frameshift mutations were more common than in-frame mutations. The red arrowhead indicates hypophosphorylated PER1 while the asterisk indicates a non-specific band. (D) Sanger sequencing of clone #2-12 identified a 27-nt (9-AA) deletion in one allele and a 14-nt (frameshifting) deletion in the second allele. (E, F) The missing 9 AAs (red) in the PAS-A deletion mutant (Allele 1) were close to the PAS-A dimerization interface (purple), as shown by AA sequence alignment and structural model. (G, H) The PAS-A deletion mutant was constitutively hypophosphorylated while wt PER2 in the mutant clone was normally phosphorylated. (I) Inhibition of PP1 by CA did not induce hyperphosphorylation of the mutant PER1. (J, K) PER1 PAS-A deletion did not disrupt interaction between CK1 and PER1 but dramatically reduced dimerization affinity. Note that some of endogenous PER1 was co-purified with transfected PER1-Luc in the last lane. NS: non-specific IgG.

Since the PER1 PAS contains two dimerization interfaces, random AA indels were also introduced in the other interface, PAS-B, to determine whether phosphorylation is similarly affected (Fig 2A). Mutant clones were selected based on altered phase and subjected to immunoblotting as described above (Fig 2B). Several AA indel clones were isolated, all of which showed severely hypophosphorylated PER1, similar to the PAS-A mutants (Fig 2B and C). As with the PAS-A mutants (#2-12), treatment of PAS-B mutants with cycloheximide (CHX) and CA failed to induce hyperphosphorylation, indicating a defect in the phosphorylation process itself. In contrast, wt PER2 in these clones remained normally hyperphosphorylated (Fig 2D, E and S2A-C). The PAS-B AA indel mutants were nonfunctional, similar to the PAS-A mutants, as bioluminescence rhythms were abolished upon *Per2* deletion (Fig 2F). However, because *Per1* mutant genes are still regulated by the *Per2*-mediated circadian feedback loop, PER1 PAS-B mutant proteins continued to oscillate in the presence of *Per2*, albeit with reduced amplitude (Fig S2D and E). As with the PAS-A mutants, a PAS-B mutant missing several AAs failed to undergo hyperphosphorylation even in transfected cells (Fig 2G). It has been shown that W448 in PAS-B is a key residue in dimer interactions, and W448E mutation would significantly disrupt this interaction ^30,38^. However, this mutation appeared less disruptive than deletions in PAS-B. When transfected, PER1 W448E was slightly less hyperphosphorylated and entered the nucleus similar to wt PER1, suggesting that the W448E PER1 mutant may be functional in vivo (Fig 2G and S2F). Both PAS-A and PAS-B mutant PER1 proteins could be hyperphosphorylated to a certain level when co-transfected with CK1 (Fig 2H and I). It has also been shown that a mutant PER lacking the entire CKBD is still hyperphosphorylated ^33^. However, this hyperphosphorylation was reduced when the PAS-A deletion was introduced into the CKBD mutant PER1 (Fig 2J).

**Fig. 2.**
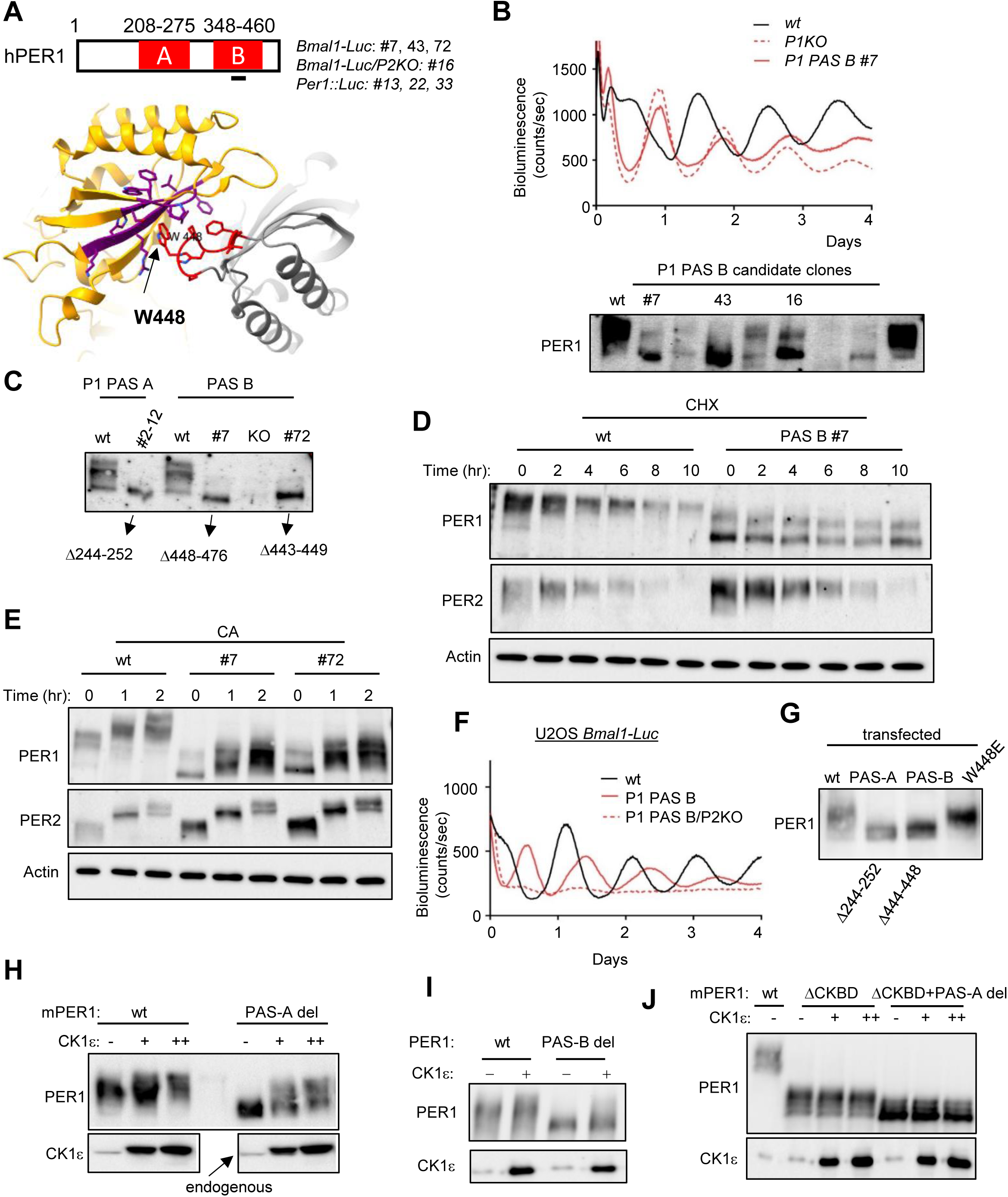
PAS-B domain in PER1 is also critical for PER phosphorylation and circadian rhythms. (A) AA indel clones in PAS-B were selected as above. W448 was targeted because it has been shown that it is one of critical residues in PAS-B dimerization interface. Indel clones were selected in three different reporter cell lines including one endogenous (knock-in) reporter *Per1::Luc*. (B-E) PER1 PAS-B deletion mutant proteins were constitutively hypophosphorylated like the PAS-A mutants. N=3 each in (B). (F) PAS-B mutant proteins were not functional. #7 (PAS-B) and #16 (PAS-B/*Per2* KO) are shown. N=3 each. (G) PER1 W448E mutant protein was more highly phosphorylated than PAS deletion mutant proteins when they were transfected. (H, I) PAS deletion mutant proteins could be hyperphosphorylated in a CK1 dose-dependent manner when they were co-transfected with CK1. ΔPAS A: 244-252, ΔPAS B: 443-449. (J) Deletion in the PAS-A domain compromised hyperphosphorylation more severely than that in CKBD. Wt PER1 and CKBD mutant PER1 could be hyperphosphorylated independent of co-transfected CK1, suggesting that endogenous CK1 is sufficient for this hyperphosphorylation. AA682-877 is missing in the CKBD mutant PER1.

We hypothesized that phosphorylation of PER2 would be similarly compromised by disruption of dimerization, given the structural homology between PER1 and PER2 (Fig 3A). To test this, we introduced random AA indels into the PER2 PAS-A domain in U2OS-*Bmal1-Luc/Per1 KO* cells as previously done for PER1. We expected that AA indel clones would be arrhythmic, similar to PER1 indel clones in the absence of the *Per2* gene, allowing for screening without molecular analysis. Indeed, arrhythmic clones had AA indels along with frameshift mutations (KO) (Fig 3A). These PER2 PAS-A mutants were hypophosphorylated relative to wt PER2, though their phosphorylation levels were higher than those observed in PER1 PAS mutants (Fig 3B). As in PER1 PAS mutants, PER2 PAS-A mutants did not exhibit excessive hyperphosphorylation upon CA treatment (Fig 3C). When a PER2 PAS-A indel clone was created in an endogenous reporter cell line, U2OS *Per1::Luc* cells, the PAS-A indel mutant was also hypophosphorylated (Fig 3D and E). The mutant PER2 showed rhythmicity in both abundance and phosphorylation, likely due to the presence of the *Per1* gene, although these rhythms were less robust than those of wt PER2 (Fig 3E). Consistent with the previous PER2 mutant, hyperphosphorylation of this mutant was not observed following the CHX and CA treatment (Fig 3F and G).

**Fig. 3.**
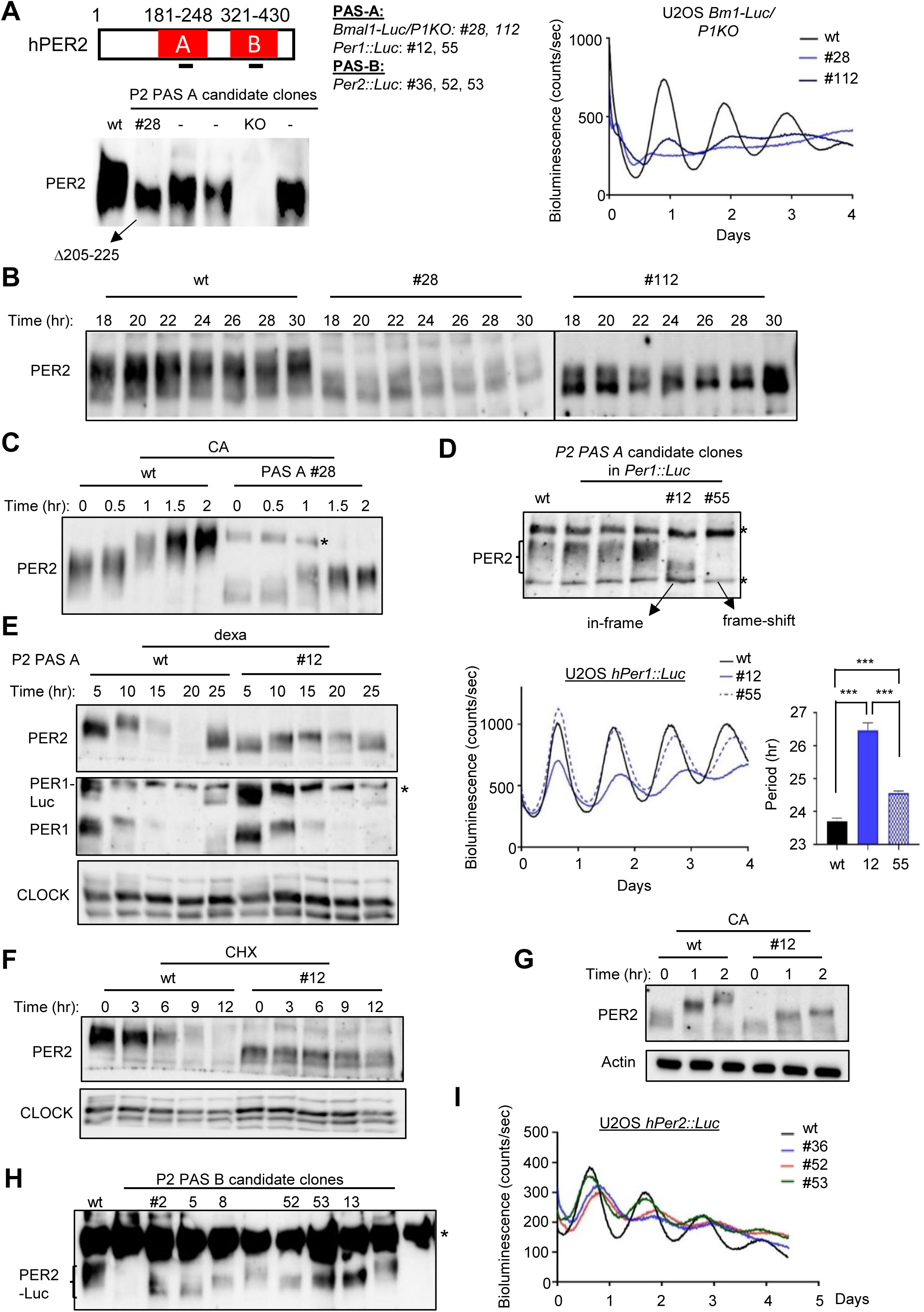
PAS domains in PER2 are also critical for PER phosphorylation and circadian rhythms. (A) AA indel clones in PER2 PAS were generated and selected as above. AA residues homologous to those targeted in PER1 PAS-A and B were targeted (see Fig 1 and 2). These clones were selected in three different reporter cell lines based on disruption of bioluminescence rhythms in the absence of PER1. N=3 each. (B) Protein blots show compromised phosphorylation of two example clones (#28, 112); two separate gels were run at the same time. The samples were harvested at indicated times after serum shock. (C) CA treatment did not induce excessive hyperphosphorylation in the mutant PER2 protein * non-specific band. (D) Hypophosphorylation was also seen in a PAS-A deletion clone #12 generated in *Per1::Luc* cells. Candidate clones were selected in a random manner. Note that phase was not shifted in #12 and #55 relative to the wt samples (bottom panel). n=3, mean+/-SEM, * p<0.05, ** p<0.01, *** p<0.001. (E) When *Per1* is intact, PER2 PAS-A deletion mutant protein oscillated in abundance although hyperphosphorylation was suppressed. (F, G) PER2 PAS-A deletion prevented hyperphosphorylation even with CHX and CA treatment. (H, I) PAS-B mutant clones were similarly hypophosphorylated as PAS-A mutants. Note that these indel mutations were made in the endogenous *Per2::Luc* reporter line. All indel mutations in the endogenous *Per2::Luc* dramatically reduced amplitude in bioluminescence rhythms consistent with (E).

When a PER2 PAS-A deletion mutant was expressed in transfected cells with or without CK1 co-transfection, it was less phosphorylated compared to wt PER2 (Fig S3A). To determine whether PER2 PAS-B also contributes to phosphotimer function, we disrupted and assessed PER2 PAS-B using the same approach. Similar to the PAS-A mutants, PAS-B indel mutants exhibited reduced level of phosphorylation compared to wt PER2 under all tested conditions (Fig 3H and S3B-D). Although PAS-B mutants appeared more phosphorylated than PAS-A mutants, they were nonfunctional (Fig S3E).

### PAS domain disruption compromises phosphorylation more than disruption in other domains

The contributions of other domains to gross phosphorylation and functionality were assessed by introducing AA indels into the PER2 degron, FASP (center of CKBD), N-terminus and C-terminus of CKBD, and the CRY-binding domain (CBD, C-terminus of PER) (Fig S4A). Disruption in the PER2 degron and CKBD-N did not affect gross phosphorylation, hyperphosphorylation, or the rhythmicity of protein phosphorylation and abundance (Fig S4B-G). However, AA indels in both domains resulted in period lengthening. In our previous study, we demonstrated that AA indels in FASP within PER1 and PER2 do not affect gross phosphorylation or protein rhythmicity but induce period shortening ^33^. These mutant proteins are likely functional in the absence of the other paralog, as a PER2 FASP deletion mutant remained rhythmic without functional *Per1* (Fig S5A-D). Interestingly, the period was further shortened in this double mutant suggesting that mutant *Per2* is semi-dominant over wt *Per1*.

We showed that a mutant PER2 missing ∼2/3 of CKBD is still hyperphosphorylated but is nonfunctional ^33^. Since this mutant PER2 still binds CK1, albeit with reduced affinity, the weak physical interaction may be sufficient for hyperphosphorylation but not for functionality. Similarly, while targeting PER1 FASP, we isolated a PER1 mutant clone missing ∼2/3 of CKBD (Fig 4A). Like the PER2 mutant, this PER1 mutant clone showed a lengthened period and was phosphorylated to a similar extent as wt PER2 (Fig 4B and C). However, it did not undergo as much excessive hyperphosphorylation as wt PER2 by CA treatment (Fig 4D). As previously suggested ^33^, a stable interaction through CKBD is probably necessary for controlled, slow phosphorylation. Given that An et al. showed the critical role of CKBD-C in PER2 hyperphosphorylation ^32^, the region was targeted by CRISPR to produce AA indels in CKBD-C in U2OS cells. Three clones were isolated, all of which were hyperphosphorylated, though to a slightly lesser extent than wt PER2. An et al. reported that hyperphosphorylated species of their CKBD-C mutant PER2 were undetectable on SDS-PAGE, which is not consistent with our results (Fig 4E-G). Our mutant clones were rhythmic with longer periods, consistent with that of the mutant mouse reported by An et al. (Fig 4E). The CKBD-C mutant PER2 proteins were rhythmic in phosphorylation and abundance comparable to wt PER2 (Fig 4F, G and S5E). However, unlike the FASP mutants., the CKBD-C mutants were nonfunctional in the absence of *Per1* (Fig S5F). In addition, the CKBD-C mutants were constitutively hypophosphorylated as reported by An et al. and less hyperphosphorylated compared to wt PER2 by CA treatment in the absence of *Per1* (Fig 4H and I). Hyperphosphorylation of the CKBD-C mutants in a wt *Per1* background appears to be mediated by heterodimerization with PER1, as CKBD-C mutant PER2 remains hypophosphorylated when PER1 PAS is compromised, similarly to when *Per1* is deleted (Fig 4J, S5G).

**Fig. 4.**
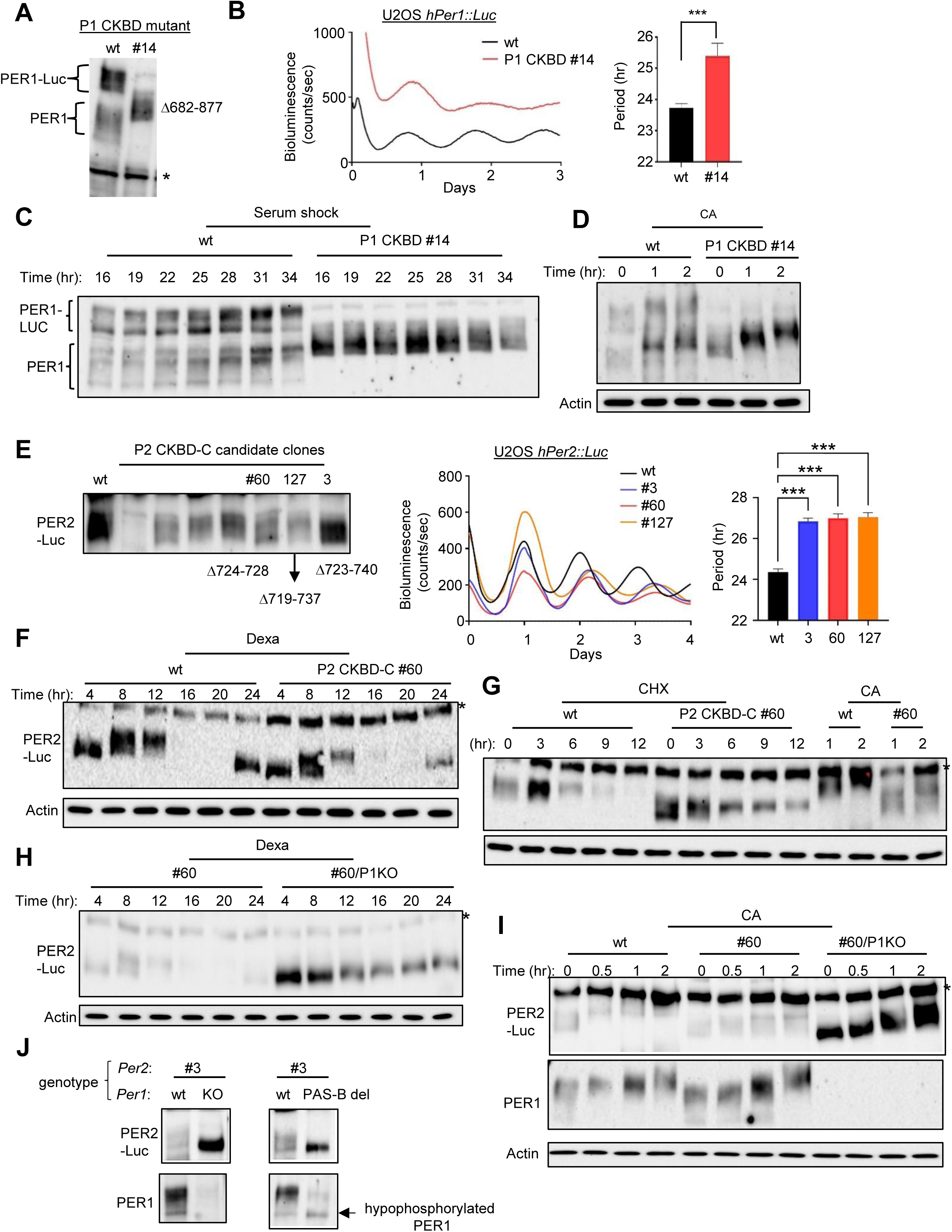
Phosphorylation is most compromised when PAS domain is disrupted compared to disruption in other domains. (A-D) A large deletion in PER1 CKBD did not compromise gross hyperphosphorylation but excessive hyperphosphorylation induced by CA treatment was truncated. Note that the maximum level of hyperphosphorylation was not altered by CA treatment for the mutant protein unlike the wt protein in (D). n=4, mean+/-SEM, * p<0.05, ** p<0.01, *** p<0.001 in (B). (E-H) CKBD-C deletion compromised hyperphosphorylation more than that in CKBD-N and FASP. Gross hyperphosphorylation of a CKBD-C deletion mutant protein was reduced, but the mutant protein oscillated in abundance and phosphorylation (F). However, these rhythms were abolished when *Per1* was deleted (H). n=3, mean+/-SEM, * p<0.05, ** p<0.01, *** p<0.001 in (E). (I) CA treatment induced less hyperphosphorylation in this mutant protein. * non-specific. (J) PER2 CKBD-C mutant proteins could be hyperphosphorylated in the presence of wt PER1, but not PER1 harboring PAS-B deletion. This suggests that heterodimerization with PER1 through the PAS domain is required for mutant PER2 hyperphosphorylation.

Finally, the CRY-binding domain (CBD) was disrupted by AA indels. Gross phosphorylation was almost intact in both PER1 and PER2 CBD mutants, suggesting that CBD is not critical for the phosphotimer (Fig S6A and B). Taken together, these data strongly suggest that the PER PAS domain is the most critical component of PER in the phosphotimer function by initiating temporally controlled, collective phosphorylation.

### Point mutations in PER PAS dramatically alter wake/sleep cycles in mouse models

Given the critical role of PER dimerization via the PAS domain in regulating PER phosphorylation kinetics, we hypothesized that disruptions in PAS-PAS dimerization would dramatically affect behavioral rhythms, including wake/sleep cycles in corresponding mutant animals. To test this hypothesis, we generated mouse models harboring SNP mutations in PAS, which would allow more nuanced functional perturbations than indel mutants and provide more relevant insights into potential circadian disorders associated with human *Per* SNPs. We targeted PER1 W448 and PER2 W419 because crystallography studies showed they are critical for PAS homodimerization, and W448E or W419E mutation disrupts dimerization ^30,38^. Furthermore, unlike PER1 PAS deletion mutants, the PER1 W448E mutant protein was hyperphosphorylated in vitro (Fig 2G), suggesting that the point mutant may be functional but could alter kinetics in vivo. Since NHEJ-mediated AA indels are common byproducts during HDR-CRISPR mutagenesis, and studying behavior of mice with such indels would complement our cell data, we also examined several AA deletion mutants ^36^. Our initial characterization of AA indel mice in both *mPer1* and *mPer2* showed results consistent with those observed in U2OS cells. Among these indel mutants, we selected one mutant for each gene: *mPer1^Δ450^*, *mPer2^Δ413-419^* (Fig S7A). These deletion mutant mice were crossed with *Per1* or *Per2* KO mice to test the functionality of these mutant proteins as above. In the C57BL6 background, both *Per1* KO and *Per2* KO exhibited robust rhythms that were nearly indistinguishable from those of wt mice (Fig 5A and S7B). As observed in U2OS cells, these indel mutant mice were rhythmic when the other paralog was present, but became arrhythmic in the absence of the paralog (Fig 5B). Circadian period was significantly lengthened in *mPer1^+/+^*; *mPer2^Δ413-419^* mutants, consistent with data in U2OS, whereas the period was slightly shortened in *mPer1^Δ450^; mPer2^+/+^* mutant mice (Fig 5C). Additionally, both deletion mutants were constitutively hypophosphorylated, regardless of the presence of the other paralog, as seen in U2OS cells (Fig 5D-F). Both deletion mutant PER proteins were also hypophosphorylated in MEFs (Fig 5G). All these data strongly support that our U2OS platform is a valid model for studying in vivo clock mechanisms.

**Fig. 5.**
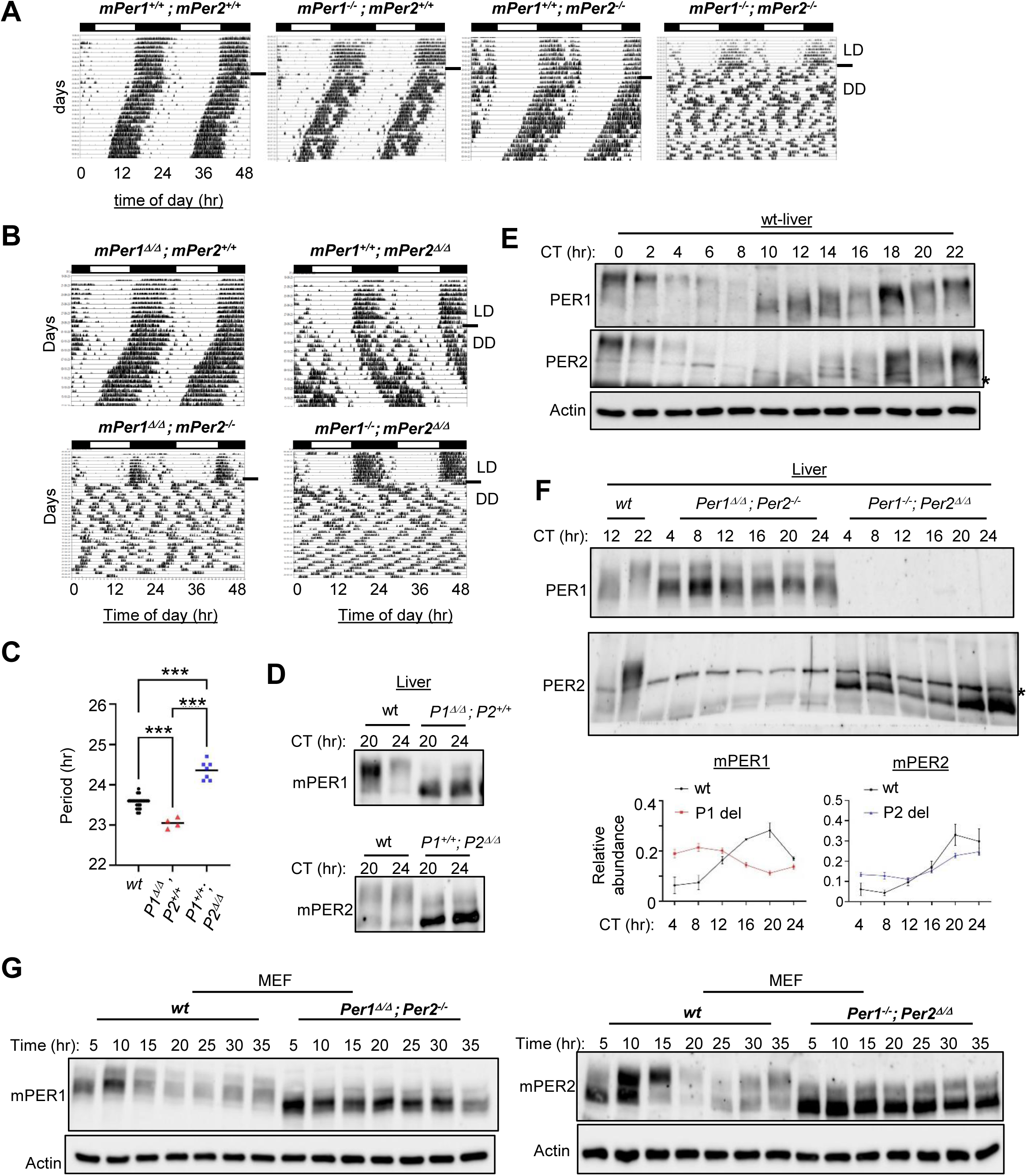
Deletion mutations in PER PAS in mouse models also induced hypophosphorylation and abolished functionality of PER. (A) *mPer1* and *mPer2* are redundant for circadian behavioral rhythms: either paralog was sufficient to sustain rhythms in locomotor activity, but double-knockout mice were arrhythmic. Representative actigraphs are shown, but the experiments included N=5 per genotype, see Fig S7B. (B, C) The deletion mutant proteins (genotype Δ/Δ) cannot sustain circadian rhythms on their own. Furthermore, in the presence of the other paralog *Per*, they seem to interfere with its function, significantly altering the circadian period. N=24 for wt, 4 for *Per1* del, 7 for *Per2* del. * p<0.05, ** p<0.01, *** p<0.001. (D-G) The deletion mutant proteins are constitutively hypophosphorylated in liver tissue in vivo and in MEFs. n=3, mean+/- SEM in (F).

Both point mutant mice were rhythmic with slightly shortened periods when the other paralog was present (Fig 6A and E). Additionally, both mutant proteins displayed robust rhythmicity in tissues, comparable to wt PER (Fig 6B). The mutant PER1 was phosphorylated to the same extent as wt PER1, whereas the mutant PER2 phosphorylation was significantly blunted. Double mutant (DM) mice were also rhythmic but had a drastically shortened period of ∼20 hrs, along with a severe phase advance (Fig 6C, E and F). Half of their daily activity occurred during the light phase, with a corresponding reduction in nighttime activity. When these mutant mice were crossed with *mPer1* KO and *mPer2* KO mice, the *mPer1^W448E^*; *mPer2* KO mice showed a similarly advanced phase and shortened rhythms. In contrast, *mPer1* KO; *mPer2^W419E^*mutant mice exhibited only a slight shortening of rhythms without advanced phase, indicating that *Per1 W448E* is mainly responsible for the dramatic phenotypes (Fig 6D and F). Both mutant proteins showed robust oscillations in MEFs, but phosphorylation of the PER2 W419E mutant was also truncated in MEFs (Fig 6G). PER rhythms in DM mice were phase-advanced by ∼8 hrs— exceeding the ∼3.5 hrs period shortening—consistent with the phase advance observed in behavioral rhythms (Fig 7A and B). Clock-controlled genes (ccgs) in DM mice exhibited a similar phase-advance (Fig 7B). While both point mutant PER proteins were similar to wt in the level of extra hyperphosphorylation when the mutant MEFs were treated with CA, this increase in phosphorylation was not observed in both deletion mutant PER proteins upon CA treatment (Fig S7C).

**Fig. 6.**
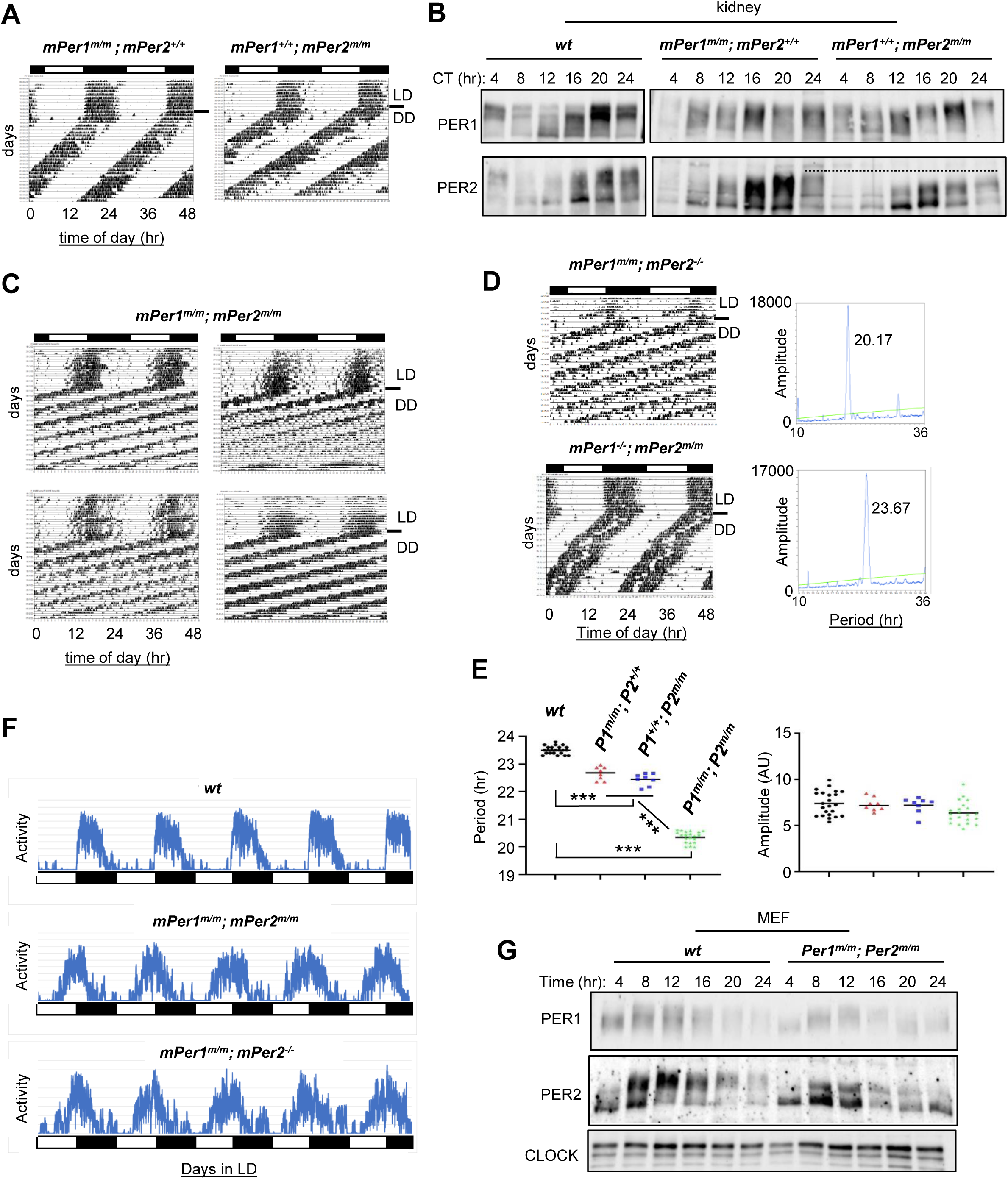
Point mutations in PER PAS dramatically alter wake/sleep cycles in mouse models. (A) Circadian rhythms are shortened in mice with single point mutations *Per1* W448E or *Per2* W419E (genotype m/m in the figures), even when the other paralog is intact. (B) Gross phosphorylation of the PER2 W419E mutant protein was truncated while the PER1 W418E mutant protein was phosphorylated as much as wt PER1. The dashed line represents a maximum phosphorylation level of wt PER2. (C-F) Double mutant mice and *mPer1*^m/m^/*mPer2* KO exhibited similarly shortened period and advanced phase; p>0.05 for period between the two mutant genotypes. N=23 for wt, 8 for *Per1* mut/*Per2* wt, 8 for *Per1* wt/*Per2* mut, 21 for DM, 7 for *Per1* mut/*Per2* KO, 8 for *Per1* KO/*Per2* mut. * p<0.05, ** p<0.01, *** p<0.001. (G) Truncated phosphorylation of the mutant mPER2 was reproduced in MEFs.

**Fig. 7.**
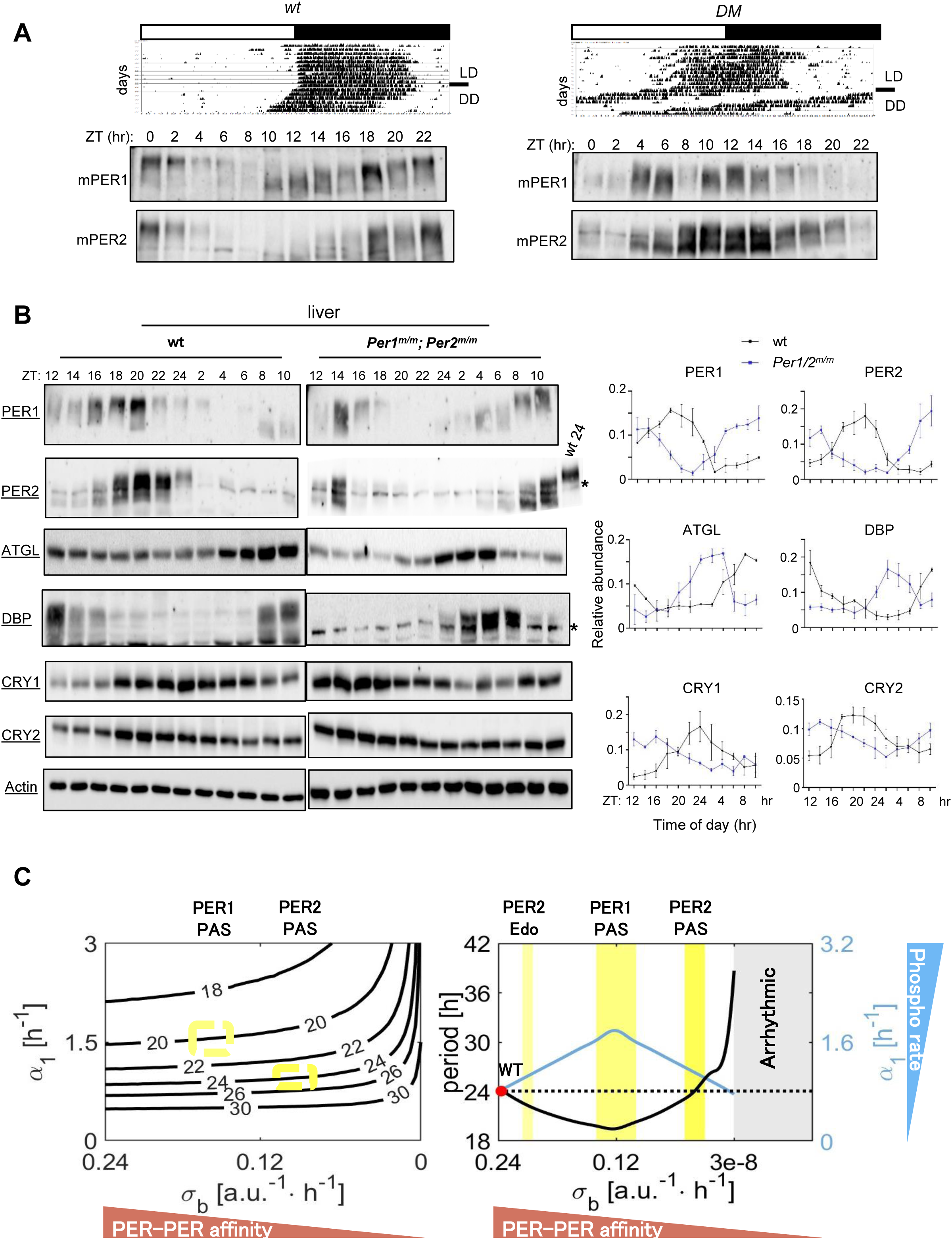
Circadian rhythms in behavior and ccgs are dramatically modulated by mutations in PER PAS domain. (A) Circadian phase was aligned between PER and behavioral rhythms in wt mice and it was similarly advanced in double-mutant mice. (B) Protein rhythms of both clock genes and ccgs were similarly advanced in double-mutant mice. When measured by JTK_Cycle, circadian phase in ccgs was advanced between 6 and 10 hrs in DM ^52^. n=3, mean+/-SEM. (C) Mathematical modeling predicted that circadian period is determined as a function of both PER:PER affinity (σ_b_) and phosphorylation rate (α_1_). As dimerization (*σ_b_*) rate decreases to a certain level (inflection point), the rate of phosphorylation (*α_1_*) increases and period decreases, which is represented by the blue and black curve, respectively, in the right graph. However, when PER:PER dimerization rate further decreases beyond the inflection point, PER phosphorylation rate decreases, which increases period length. Based on our model, we propose that disruption in PER1 dimerization by W448E changes the default value (wt) to near the inflection point (medium yellow) while the PER2 mutation disrupts dimerization more severely to the bright yellow zone. The *Per2^Edo^* mutation (I324N) would disrupt dimer modestly (light yellow). When the rate of dimerization (*σ_b_*) is less than the threshold ( ∼3 ⋅ 10^−8^) as in our deletion mutants, oscillations vanish, as represented by the gray shaded area. As the rate of dimerization (*σ_b_*) and the rate of phosphorylation (*α_1_*) vary, the period of circadian rhythms is altered in the left graph. Solid black curves represent the values of parameters *σ_b_* and *α_1_* that generate oscillations of period (in hrs) as labeled. PER1 mutant and PER2 mutant exhibited rhythms of period of approximately 20 hrs and 24 hrs respectively, with its region indicated on the graph.

### Circadian timing cues are encoded in PER dimerization-mediated trans-phosphorylation

Our data demonstrated that PER phosphorylation absolutely requires dimerization, which confirms our previous modeling prediction of the circadian feedback loop. Our previous mathematical model suggested that dimerization or multimerization would be required for collective phosphorylation of individual molecules which leads to synchronous nuclear entry of the inhibitor complex after time delay in a non-discrete manner ^35^. Since nuclear entry-mediating hyperphosphorylation is a trans-acting process catalyzed by CK1δ/ε in PER dimers or multimers, this would be facilitated only when PER levels surpass a certain threshold and increases dramatically thereafter. Therefore, this model generates a robust feedback inhibition within a narrow time range after a time delay in feedback inhibition. As demonstrated in our study and others ^19,21,27,32^, the kinetics of PER phosphorylation is also affected by PER-CK1 affinity. However, dimerization or multimerization of PER is required for functional PER:CK1 interaction. Subtle compromise in PER:CK1 interaction caused by mutations and deletions in FASP (central region of CKBD) induces accelerated phosphorylation kinetics and consequently period shortening ^21,33^. Similarly, in our current study, weakening in PAS-PAS interaction led to period shortening, more dramatic than the FASP mutation. These data are, however, counterintuitive because mutations resulting in reduction of PER dimerization would lengthen the period due to slower accumulation of PER dimers. To investigate this conundrum, we constructed a mathematical model that explicitly describes the dimerization and subsequent phosphorylation of PER (Fig 7C).

Consistent with our intuition, the period monotonically increases as the rate of PER dimerization, σ_b_, decreases, which is opposite of the observed period shortening. On the other hand, our model indicates that the period decreases as the rate of phosphorylation, α_1_ increases, which is consistent with previous studies ^21,33^. These data suggest that the observed period phenotypes are probably due to an integrative change between σ_b_ and α_1_ (Fig 7C). In other words, the mutations that weaken PER dimerization somehow results in an increase in the rate of phosphorylation, compensating for the period lengthening caused by weakened PER:PER interaction. Since trans-phosphorylation would depend on the probability of physical interactions between CK1 and distant phosphorylation sites, weakening of PER:PER interaction would increase the flexibility of these complexes, thereby enhancing the likelihood of interaction and leading to an acceleration of the phosphorylation kinetics. A normal circadian period can be maintained when PAS-PAS affinity is only slightly compromised. With significantly reduced affinity, trans-phosphorylation is enhanced due to increased interaction between the kinase and the substrate, analogous to the acceleration of reaction rates between two free molecules with increasing temperature. However, with a further reduction in affinity, PER dimers would eventually become too loose to maintain functional proximity between two monomers, resulting in constitutive hypophosphorylation and arrhythmicity. While PER1 W448E causes a dramatic shortening of the period, PER2 W419E does not significantly affect the period. One would expect that phosphorylation of the mutant PER1 would be more compromised than that of the mutant PER2. However, in both tissues and MEFs, gross phosphorylation of PER1 remains intact, while that of the mutant PER2 is significantly truncated. Our mathematical model theoretically explains this highly counterintuitive data. We propose that the mutant PER2 dimer affinity is more compromised than that of the PER1 mutant dimer, yet still within a permissible range for sustained rhythms. However, in this case, excessively weakened dimer affinity may cancel out the increased reaction rate between CK1 and PER, resulting in a wt-like period and truncated hyperphosphorylation. This model can explain the shortened period (∼1.5 hrs) of the previous *Per2* PAS mutant mouse, *Per2^Edo^* ^39^. The *Per2* mutant mouse was identified in a forward genetic screening and turned out to have a point mutation between PAS-A and PAS-B. This mutation does not disrupt PER2 dimerization as severely as W419E, but would compromise dimer affinity ^40^. We hypothesize the existence of an inflection point in PAS dimer affinity at which the increased reaction rate is counterbalanced by the weakened dimer affinity, ultimately stabilizing the period length.

## Discussion

Our extensive mutational analysis in PER strongly suggests that trans-phosphorylation through PAS dimerization predominates over cis-phosphorylation and drives the phosphotimer to generate precise temporal cues for critical events such as nuclear translocation and feedback inhibition ^35^. Based on available mutant cell and mouse models, circadian phase and period can be determined and modulated by mutations in any essential clock genes ^33,41,42^. Because all the components are intricately interconnected, the effects of certain mutations may be indirect through affecting more critical components. We believe that key circadian parameters such as phase and period are directly modulated by mutations in the components of the phosphotimer, whereas mutations in other clock genes influence these parameters indirectly by altering expression and activity of the key components. Consistently, circadian period and phase are most dramatically modulated by mutations in *CK1* and *Per* genes without compromising robustness of the oscillator ^21,26,33,43–45^. A mouse model harboring the *CK1 tau* mutation shows a dramatically shortened period of ∼20 hrs, similar to our PAS mutant mice ^43^. However, there is a striking difference between these two period shortenings. In *tau* mice, there is little phase advance or activity during daytime in LD entrainment, suggesting that active or nocturnal phase was asymmetrically more compressed ^43^. However, in our PAS mutant mice, diurnal phase is more compressed, leading to a dramatic advance of activity onset to nearly the middle of the day. This is particularly striking because mouse activity is strongly suppressed by light, regardless of circadian phase, through a behavior known as masking ^46^. Masking is considered dominant over the circadian clock, likely serving as an innate defensive mechanism against predation in nocturnal species such as mice. However, in our mutant mice, the circadian clock seems to override the masking behavior, and this suggests that the temporal niche in relation to the light cycle may not be as strict as previously assumed. As shown in this study, any disruptions in PER dimerization significantly impact behavioral rhythms, including wake/sleep cycles and overall physiology in corresponding mutant animals, as evidenced by dramatic shifts in both behavioral and ccg rhythms. It has been also shown that disruption of the dimerization of a PER analog, FRQ, in the model filamentous fungus, *Neurospora crassa*, abolishes circadian rhythms ^47^, which suggests that dimerization of this negative element is a conserved mechanism for the phosphotimer in the eukaryotic circadian clock machinery. Since there are many human missense SNPs in and around PAS according to the SNP database, and some of them may significantly disrupt PAS dimerization, it is likely that certain human circadian sleep disorders are associated with these mutations and could be more severe than the FASP syndrome. However, these severe sleep disorders would be manifested only when one of the two paralogs is inactivated by a frameshift or nonsense mutations, as PAS mutations are not as dominant as FASP mutations.

Our trans-phosphorylation mechanism fills major gaps in our understanding of the molecular clock. As demonstrated in our models, our findings can explain how the phosphotimer is integrated into the circadian feedback loop, generates time cues, and delays before the initiation of feedback in a biologically compatible, non-discrete manner. Our model also explains how the robustness of the pacemaker PER molecular rhythms is collectively generated. The collective PER phosphorylation mechanism generates an ideal kinetics curve for a self-sustaining negative feedback loop previously conceptualized by Goodwin even before the identification of any clock components ^48^. Furthermore, our in silico model can predict how much period and phase can be shifted by specific mutations in PAS because their relative disruption in PAS dimerization can be measured against three reference PAS mutations in vitro.

## Materials and Methods

### Experiments in mice and MEFs

#### Generation of mutant mice and genotyping

All mice were maintained in a climate-controlled room and used according to the Florida State University Animal Use Committee’s guidelines. All experiments involving animals were performed according to approved protocols by FSU ACUC, protocol number: 202200000021. All methods are reported in accordance with ARRIVE guidelines. Generation of mutant mice with point mutations and deletions has been described in our previous study ^36^. We used about equal numbers of male and female mice. Sex differences in behavioral rhythms and cellular clock rhythms are very subtle to insignificant ^49^. C57BL/6J (#000664) wt mice were purchased from the Jackson Lab.

Genomic DNA was extracted from mouse ear tissue. Briefly, tissue was digested in extraction buffer (100 mM Tris-Cl (pH8.5), 5 mM EDTA (pH8.0), 0.2% SDS, 200 mM NaCl, 0.4 U/ml Protease K, and 300 µg/ml RNase A) for 2-4 hours at 55 ℃. The digested tissue was centrifuged in a mini Eppendorf centrifuge at 12,000 rpm for 5 minutes, the supernatant was transferred to a new tube, and 0.8 volume of isopropanol was added to precipitate DNA. The DNA was pelleted and separated from solution via centrifugation and washed twice with 70% ethanol, dried at RT, and dissolved in 100 µl of TE buffer (pH8.0). One or two µl of the extracted DNA was used for genotyping PCR (NEB, Phusion polymerase). The PCR amplicons were analyzed by a 1.5% agarose gel and/or a 12% TBE polyacrylamide gel. PCR conditions are as follows.

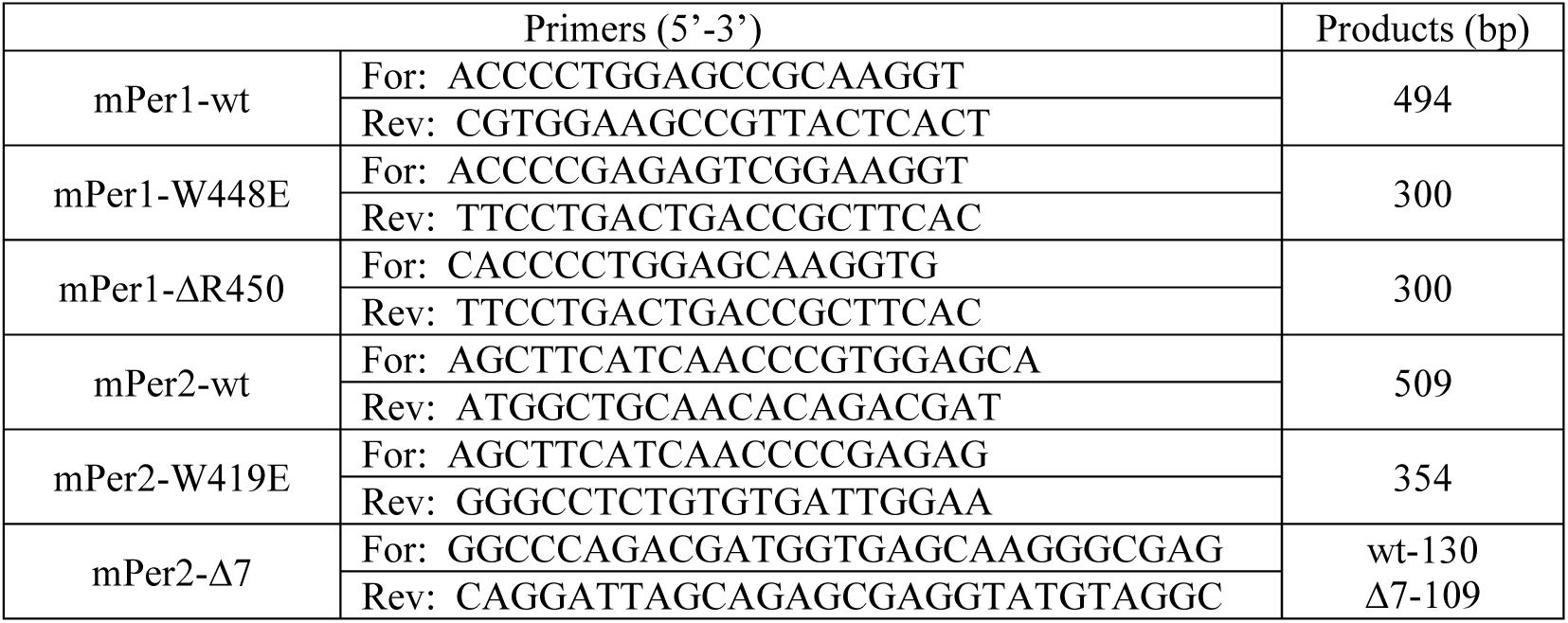

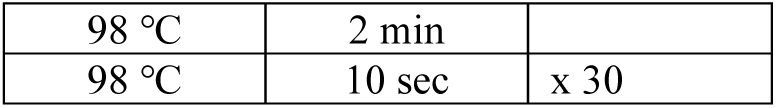
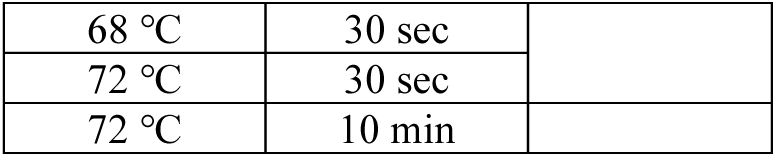

#### Analysis of circadian behavioral rhythms and tissue collection

All mice were individually housed in wheel-running cages, with free access to food and water. Wheel running activity was recorded and analyzed using ClockLab (Actimetrics, Wilmette, IL). All animals between 3 and 10 months old were initially placed in a 12h light:12h dark (LD) cycle, for at least 14 days. Mice were then transferred to constant darkness (DD), for more than four weeks, to measure clock-driven rhythms. Activity data were recorded in 5 min bins using Scurry Activity Monitoring Software (Lafayette Instrument, Lafayette, IN). Period and activity levels were calculated using ClockLab (Version 6; Actimetrics, Wilmette, IL). See the Source Data File for all raw actograms. In Fig 5 and 6, mouse tissues were collected on the first day in DD while in Fig 7, tissues were collected in LD.

#### Mouse Embryonic Fibroblasts (MEFs)

MEFs were prepared from embryos isolated from pregnant female mice, 13 days post-coitum. Embryos were removed, finely minced and treated with 0.25% trypsin and incubated at 37 °C for 30 minutes. The mixture was passed through a fine 100µm membrane to remove debris, and the resulting cells were maintained at 37 °C, 5% CO_2_ in Dulbecco’s Modified Eagle Medium (DMEM), supplemented with 10% fetal bovine serum (FBS). Both male and female embryos were used. Samples were collected at indicated times after 2 hr serum shock with 50% horse serum.

### Cell Lines

U2OS (ATCC HTB-96) cells were obtained from the commercial vendor ATCC. Transient expression of proteins by transfection was done in U2OS cells. The U2OS-*Bmal1-luc* and *Bmal1-luc*/*Per2* KO (*Per2*-E5-2) cell lines were described previously ^50^. U2OS-*Bmal1-luc*/*Per1* KO and U2OS-*Bmal1-luc*/*Per1/2* KO clones were selected from our previous study ^50^. In the study we showed that *Per1* can be efficiently knocked out when exon 6 was targeted by CRISPR ^50^. The *Per* double KO clone was generated by knocking out *Per1* in the *Per2*-E5-2 clone. The endogenous *Per1::Luc* (H10) and *Per2::Luc* (LH1) clones were described previously ^33^. All cells were maintained at 37 °C, 5% CO_2_ in DMEM, supplemented with 10% FBS. See Table S1 for cell lines generated in the current study.

### Bioluminescence recording

Cells were plated into 24-well plates or 35 mm dishes to be approximately 90% confluent 24 hours prior to the start of the experiment. Immediately before the start of the experiment, cells were given a two-hour serum shock with 50% horse serum in DMEM or 0.2μM dexamethasone in DMEM, washed with phosphate-buffered saline (PBS) and fresh DMEM supplemented with 1% FBS, 7.5mM sodium bicarbonate, 10 mM HEPES, 25 U/ml penicillin and 25 μg/ml streptomycin. Also 0.1mM luciferin was added. The plates were sealed with cellophane tape and the dishes with cover glass with vacuum grease, and placed into a LumiCycle 96 or 32 (Actimetrics, Wilmette, IL). For all bioluminescence experiments, the results were reproduced in at least two independent experiments. Real-time levels, period, and phase of the bioluminescence rhythms were evaluated using the LumiCycle software (Actimetrics).

### Drug treatments in cells

Cells were seeded in 60-mm dishes for immunoblots or 24-well plates for bioluminescence monitoring to be 90% confluent 24 hours prior to the experiments with drug treatments. For cycloheximide (CHX) treatment, 8 μg/ml was added to cells and cells were collected at specified times after the treatment or placed into a LumiCycle 96 for bioluminescence monitoring. For CHX washout experiments, cells were treated with CHX for 12 hrs followed by the normal DMEM, and cells were harvested at the indicated times after the washout. 20 nM Calyculin A (CA, EMD chemicals) were used for cells.

### Plasmids, transfection and Microscopy

*mPer1-Luc* plasmid in Fig 1K was generated by cloning *mPer1-Luc* into pAdTrack-CMV vector (Addgene #16405). *pcDNA-Per1*, *Per2, CK1*ε and *CK1δ* plasmids were described previously ^13^. The *Per1-PAS-A* deletion mutation in Fig 1K was generated by deleting the same 9 AA in this *pcDNA-Per1* plasmid. *Per1-PAS-B* deletion, W448E and ΔCKBD mutants were also generated in this plasmid. For Fig S1F, inducible *mPer1-mCherry* and *mPer1 PAS-A* deletion-Venus transgenes were used. These inducible transgenes were generated by replacing *mPer2-Venus* in the previously described inducible *tetO-Per2-Venus*; *CMV-rtTA*-pAdTrack plasmid with these transgenes ^35^. The inducible plasmids were cotransfected into U2OS cells. Induction with 1 μg/ml doxycycline was initiated 24 hrs later after transfection. The images were taken continuously for 21 hrs after induction using Andor revolution spinning disk laser confocal microscope with a live-cell chamber maintained at 37 °C and 5% CO₂. For Fig S2F, *Per1* mutations were generated in the inducible *mPer1-Venus* plasmid described above. The images were taken 24 hrs later after induction. For Fig S3A, the *mPer2 PAS-A* mutant was generated in the pcDNA plasmid described above. In Fig 2G, an equal amount (0.5 ug) of *wt Per1* and mutant *Per1* plasmids were transfected into U2OS cells. In Fig 1H-J, an equal amount of *mPer1* plasmids and 0, 10 or 50 ng *CK1ε* plasmid were used. In Fig S3A, 0.5 μg *Per2* plasmids were used with 0, 10 or 50 *CK1ε* or 50 ng *CK1δ* plasmid were used. All transfection experiments were done using jetOPTIMUS (Polyplus, NY, NY).

### Prediction of protein folding

To investigate the effects of mutations on dimerization, protein folding of wild-type and mutant dimers was predicted for mPER1 PAS (wild-type: PDB ID [4DJ2], residues 191–502; mutant: mPER1[197–502] W448E) and mPER2 PAS (wild-type: PDB ID [3GDI], residues 170–473; mutant: mPER2[170–473] W419E). Structure predictions were performed using ColabFold, a user-friendly implementation of AlphaFold. For each construct, five structural models were generated and ranked based on confidence scores. Predicted Local Distance Difference Test (pLDDT) scores and Predicted Aligned Error (PAE) matrices were computed to assess model confidence. Only the highest-ranked model for each construct was selected for downstream analysis. The resulting 3D structures were visualized, and the dimerization interface was examined using UCSF ChimeraX.

### Immunoblotting and Immunoprecipitation

The cells in 6 cm dishes were harvested and flash-frozen on dry ice. Protein extraction and immunoblotting were performed as previously described ^51^. Briefly, cells were homogenized at 4°C in 70 μl extraction buffer (EB) (0.4M NaCl, 20mM HEPES (pH 7.5), 1mM EDTA, 5mM NaF, 1 mM dithiothreitol, 0.3% Triton X-100, 5% glycerol, 0.25mM phenylmethylsulfonyl fluoride, 10mg of aprotinin per ml, 5mg of leupeptin per ml, 1mg of pepstatin A per ml). Homogenates were cleared by centrifugation 12 min, 12,000g at 4°C. Supernatants were mixed with 2x sample buffer and boiled. Proteins were separated by electrophoresis through SDS polyacrylamide gels and then transferred to nitrocellulose membranes. Membranes were blocked with 5% non-fat dry milk in TBS-0.05% Tween-20 (TBST), incubated with primary antibodies overnight followed by incubation with secondary antibodies for 1 hr. The blots were developed using an enhanced chemiluminescence substrate (WestFemto, ThermoFisher Scientific).

Antibodies to clock proteins were described previously ^17,19,50^. hP2-GP49 (hPER2), GP58 (mPER2), GP62 (PER1), CLK-1-GP, BM1-2-GP, C1-GP (CRY1), C2-GP (CRY2), CK1δ-GP and CK1ε-GP antibodies were used at 1:1,000 dilution in 5% milk–TBST solution. Anti-DBP polyclonal antibodies were generated guinea pigs as described previously ^17^. GP38 was used. Anti-ATGL antibody was purchased from Cell Signaling, #2138 (Danvers, MA01923). Rabbit anti-ACTIN antibody (Sigma, A5060) was used at 1:2,000.

Immunoprecipitation was performed as described previously ^17^. Briefly, protein extracts from cells harvested from 10 cm dishes were prepared as described above. 10% of the initial protein extract was saved for the input. 20μL Protein G Sepharose 4 Fast Flow beads (GE Healthcare) per reaction was equilibrated with 500μL of EB for 15 minutes on a rotating wheel. The beads were centrifuged at 3000 rpm for 15 seconds and the supernatant was removed. This wash step was repeated three additional times. After the final wash, two volumes of EB were added to the beads. To pre-clear the extracts, 10μL of the equilibrated bead solution were added to the extracts and incubated for 30 minutes on a rotating wheel at 4°C. The samples were centrifuged at 12,000 rpm for 5 minutes at 4°C, and then the pre-cleared extract was transferred to a fresh tube. Then 0.1μg of affinity-purified antibody and 10 μl of the equilibrated beads were added to the tube. This mixture was incubated at 4°C on a rotating wheel for four hours. The tubes were centrifuged at 3000 rpm for fifteen seconds, and the supernatant was removed. 1mL of EB was added to the tube and incubated on a rotating wheel at 4°C for twenty minutes. The samples were centrifuged at 3000 rpm for 15 seconds, and the supernatant was removed. This step was repeated three more times to completely wash the beads. After the final wash, the majority of the EB was removed and 30μL of 1X sample buffer was added. The samples were boiled at 95°C for 3 minutes.

### Data analysis and data reproducibility

All statistical analyses involving multiple groups were performed using GraphPad Prism. Group comparisons were conducted by ANOVA to assess overall differences among multiple groups. When the ANOVA indicated a significant effect (p < 0.05), Tukey’s post hoc test was applied to determine pairwise differences between group means. A p-value of less than 0.05 was considered statistically significant. For analysis involving only two groups, Student’s t-test was used. *p<0.05, **p<0.01, ***p<0.001, ****p<0.0001. See the Source Data file for all statistics.

#### JTK_CYCLE

To identify rhythmic patterns in gene expression, we applied the JTK_CYCLE algorithm, a non-parametric method optimized for detecting periodic signals in time-series data ^52^. The analysis was performed using the JTK_CYCLE implementation in R ^52^, specifying a period range of 12– 28 hours to reflect expected circadian-like oscillations. Input data consisted of normalized expression values sampled at either 2h or 4h time intervals. The algorithm outputs adjusted p-values (BH.Q), period, phase (lag), and amplitude for each tested gene. Rhythmicity was defined as having a Benjamini-Hochberg adjusted p-value (BH.Q) less than 0.05.

All LumiCycle and immunoblot results have been reproduced in at least two independent experiments and representative results are shown.

### Mathematical modeling

We developed a mathematical model to illustrate the role of PER dimerization in the circadian clock, which is a system of ordinary differential equations (ODEs) consisting of 10 variables. Each variable represents the concentration of a component in the PER dimerization network, where *M* represents *Per* mRNA, *P* and *C* represent the monomer and dimer PER respectively, *C*_1_, ⋯, *C*_4_ represent the cytoplasmic PER species in each phosphorylation step, *C_N_* represents the nuclear PER that are not bound to CLOCK:BMAL1 (activator complex), *B* represents CLOCK:BMAL1 that are not bound to PER:CRY, and *I* represents the inactive complex of CLOCK:BMAL1 bound to PER:CRY. Note that CRY is not explicitly considered as a variable in our model. A system of ODEs is given in the following. Default values of each parameter are given in Table 1.

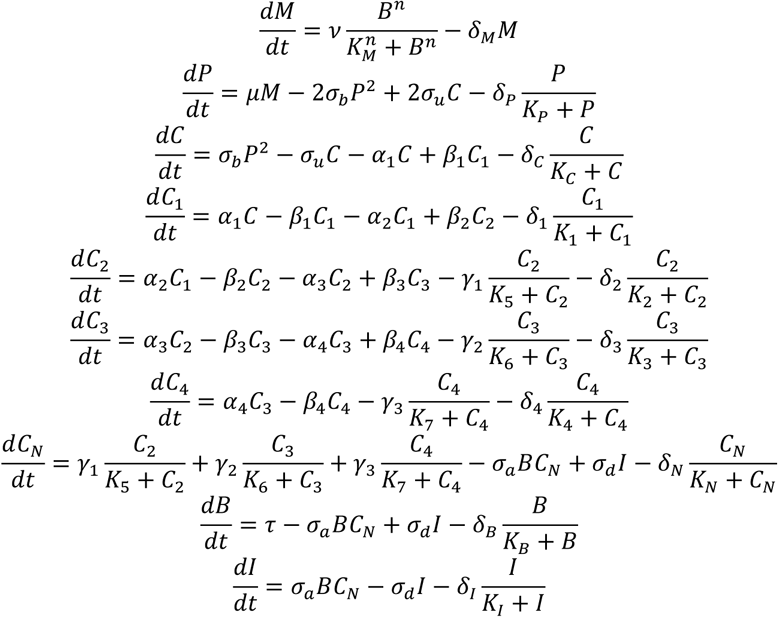

**Table 1.**

| Parameter | Description | Default | Unit |
| --- | --- | --- | --- |
| $v$ | Rate of transcription | 4.6385 | $a.u. \cdot h^{-1}$ |
| $\alpha_1, \alpha_2, \alpha_3, \alpha_4$ | Rate of phosphorylation | 0.8497, 6.8854, 14.1643, 19.2971 | $h^{-1}$ |
| $\beta_1, \beta_2, \beta_3, \beta_4$ | Rate of dephosphorylation | 0.2207, 6.5168, 7.2065, 5.8500 | $h^{-1}$ |
| $\gamma_1, \gamma_2, \gamma_3$ | Rate of nuclear transport | 32.7189, 35.9934, 38.7716 | $a.u. \cdot h^{-1}$ |
| $\mu$ | Rate of translation | 3.3407 | $h^{-1}$ |
| $\sigma_b$ | Rate of binding (dimerization) | 0.2309 | $a.u.^{-1} \cdot h^{-1}$ |
| $\sigma_u$ | Rate of unbinding (de-dimerization) | 0.5853 | $h^{-1}$ |
| $\sigma_a$ | Rate of binding (inactive complex) | 0.9338 | $a.u.^{-1} \cdot h^{-1}$ |
| $\sigma_d$ | Rate of unbinding (inactive complex) | 2.8143 | $h^{-1}$ |
| $\delta_M$ | Rate of degradation of *Per* mRNA | 0.1455 | $h^{-1}$ |
| $\delta_P, \delta_C, \delta_N, \delta_B, \delta_I$ | Rate of degradation | 3.5153, 4.2148, 0.8163, 1.3585, 7.9811 | $a.u. \cdot h^{-1}$ |
| $\delta_1, \delta_2, \delta_3, \delta_4$ | Rate of degradation of phosphorylated dimer | 6.7424, 2.1810, 1.5452, 1.0121 | $a.u. \cdot h^{-1}$ |
| $K_M$ | Threshold concentration for transcription | 10.5 | $a.u.$ |
| $K_P, K_C, K_N, K_B, K_I$ | MM constant for degradation | 1.7, 1.1, 15.2, 2.3, 3.3 | $a.u.$ |
| $K_1, K_2, K_3, K_4$ | MM constant for degradation | 0.4, 5.3, 0.4, 2 | $a.u.$ |
| $K_5, K_6, K_7$ | MM constant for nuclear transport | 2.1, 4.8, 6.5 | $a.u.$ |
| $\tau$ | Rate of production of CLOCK:BMAL1 | 5.53949 | $a.u. \cdot h^{-1}$ |
| $n$ | Hill coefficient for repression | 4 | . |

See *SI Appendix* for more materials and methods.

## RESOURCE AVAILABILITY

## CORRESPONDENCE

Requests for further information and resources should be directed to and will be fulfilled by the lead contact

## MATERIALS AVAILABILITY

Plasmids, antibodies, Matlab codes and mutant mice will be available from the corresponding author upon request.

## DATA AVIALBILITY

Uncropped immunoblot scans, all actograms and numerical source data supporting the findings of this study are provided as Source Data files. All other data supporting the findings of this study are available from the corresponding author on reasonable request. Source data are provided with this paper.

### ACKNOWLEDGMENTS

We thank Dennis Chang for assistance with manuscript preparation, the UTSW Transgenic core facility for generating mutant mice and FSU FACS facility for single cell sorting.

This work was supported by NIH grants GM147340, GM131283 and Ed and Ethel Moore AD Research Program 24A04 (C.L.), University of Cincinnati College of Medicine Bridge Funding (C.I.H.).

## AUTHOR CONTRIBUTIONS

Conceptualization, C.L., K.L., C.H. and S.L.; methodology, K.L., J.P. and J.L.; investigation, K.L., J.P., J.L. and J. H..; data analysis and modeling, C.L., K.L., J.L. and J.P..; supervision, C.L., C.H. and S.L.; writing original draft, C.L., K.L., C.H. and S.L.; and writing– review and editing, C.L., K.L., J.P., J.L., C.H. and S.L.

## DECLARATION OF INTERESTS

The authors declare no competing interests.

## Notes

### Competing Interest Statement

The authors have declared no competing interest.

