## Supplementary Information for "Circadian phosphotimer generates time cues through PER dimerization-mediated trans-phosphorylation"

Fig S1-8
Table S1: Cell lines used in this study

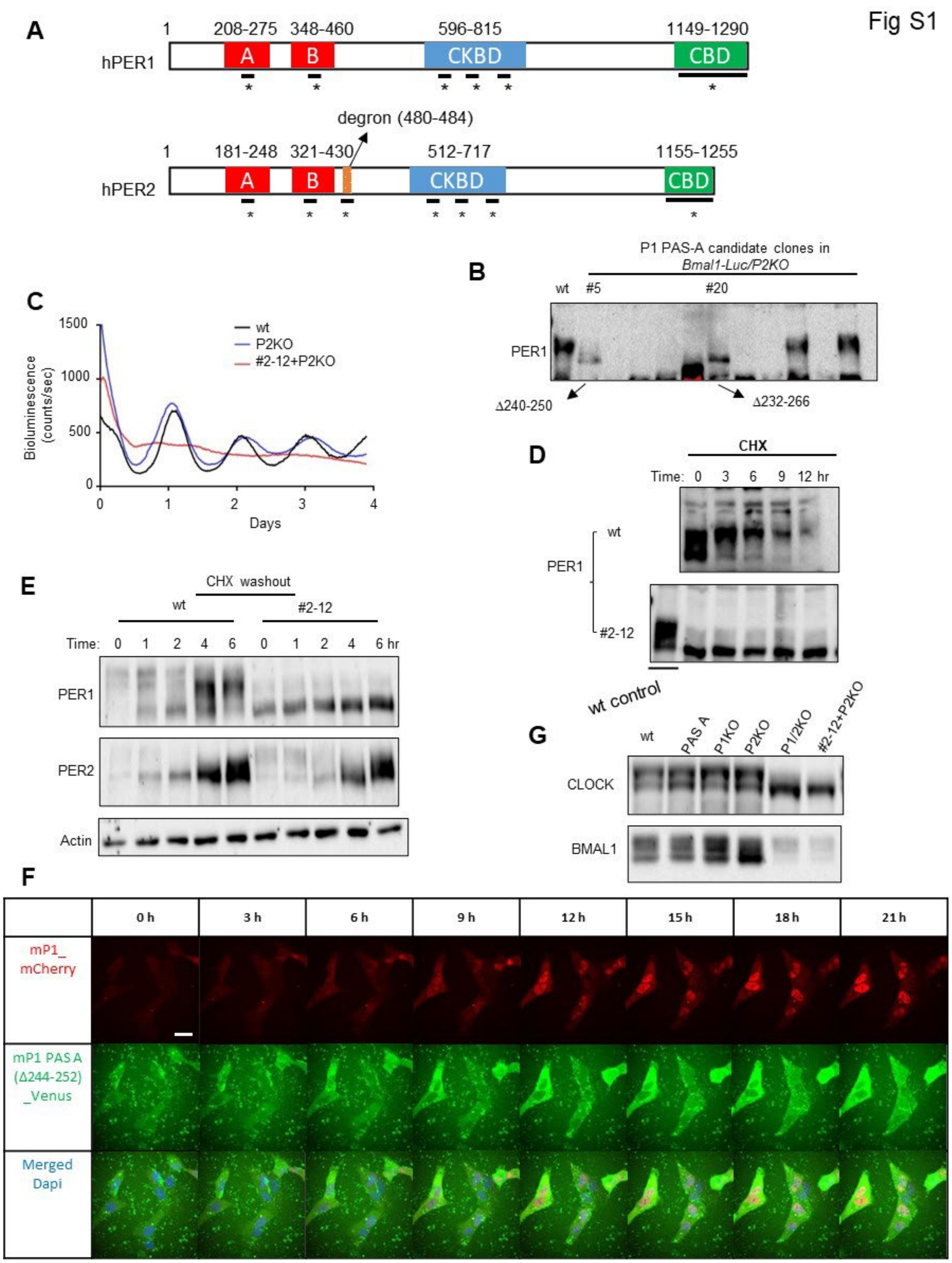

**Fig S1. PAS-A domain in PER1 is critical for PER1 phosphorylation and circadian functionality, related to Fig 1.**

(A) Schematic diagram of PER1 and PER2 domains and motifs. Asterisks represent target regions for random AA indels. Mutant clones with AA indels in FASP region have been also generated and reported in our previous study [1]**.** (B) AA indel clones in *Bmal1-Luc*/*Per2* KO line. (C) The mutant PER1 protein in clone #2-12 is not functional. *Per2* was deleted by targeting exon 5 using our previous confirmed sgRNA [2]. The single clone was selected by FACS and confirmed by immunoblotting and sequencing. The *Per2* KO clone (E5-2) has been described previously [2]. (D and E) The PER1 PAS-A mutant is constitutively hypophosphorylated. De novo PER was measured by CHX treatment for 12 hrs followed by washout in (E). (F) The mutant PER1 is always cytoplasmic. Inducible transgenic wt and PAS-A mutant PER1 labeled with mCherry and Venus, respectively, were coexpressed in U2OS cells by transfection and visualized by confocal microscopy. The scale bar indicates 40 μm. (G) CLOCK phosphorylation is suppressed in both *Per1/2* double-KO and the #2-12 clone with indel mutant *Per1* combined with *Per2* KO.

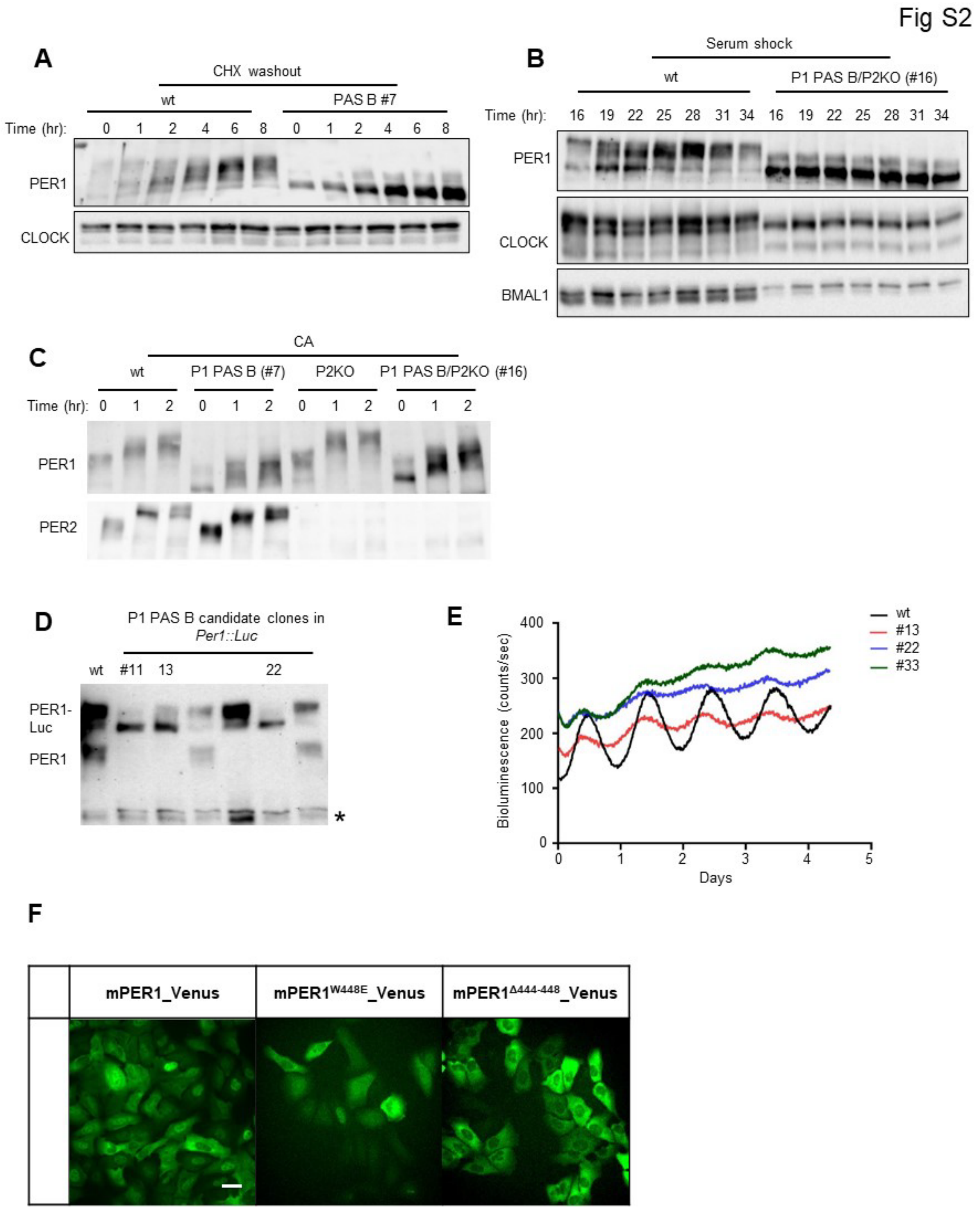

**Fig S2. PAS-B domain in PER1 is also critical for PER1 phosphorylation and circadian functionality, related to Fig 2.**

(A-E) PER1 PAS-B mutant proteins are constitutively hypophosphorylated. Note that CLOCK is hypophosphorylated in a PAS-B mutant clone combined with *Per2* KO in (B). (F) Like the PAS-A mutant PER1, PAS-B mutant PER1 is also cytoplasmic. The scale bar indicates 20 μm.

Fig S3

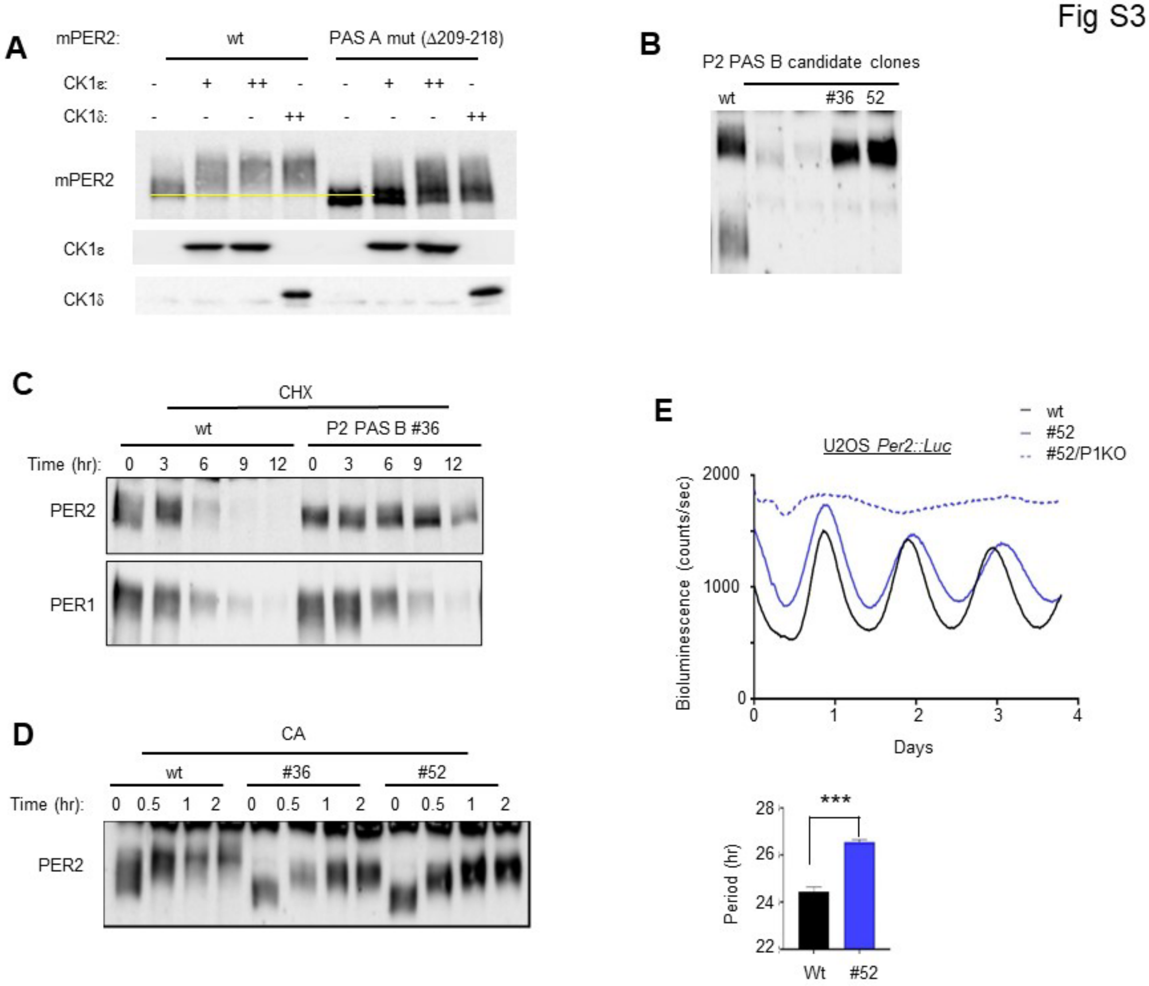

**Fig S3. AA indels in PER2 PAS domain disrupt phosphorylation and functionality of PER2, related to Fig 3.** (A) A PER2 PAS-A deletion mutant is hypophosphorylated compared to wt PER2 when they are transiently expressed. The yellow line indicates bottom of wt PER2 band. (B-D) PAS-B deletion mutants are hypophosphorylated compared to wt PER2 in various conditions, similar to PER2 PAS-A mutants. Note that the PAS-B mutant is not hyperphosphorylated and is more stable than wt PER2 after CHX treatment (C). (E) Although PER2 PAS mutants are apparently more phosphorylated than those of PER1, they are not functional and do not sustain circadian rhythms in reporter cells lacking *Per1*. n=3, mean+/-SEM, * $p<0.05$, ** $p<0.01$, *** $p<0.001$.

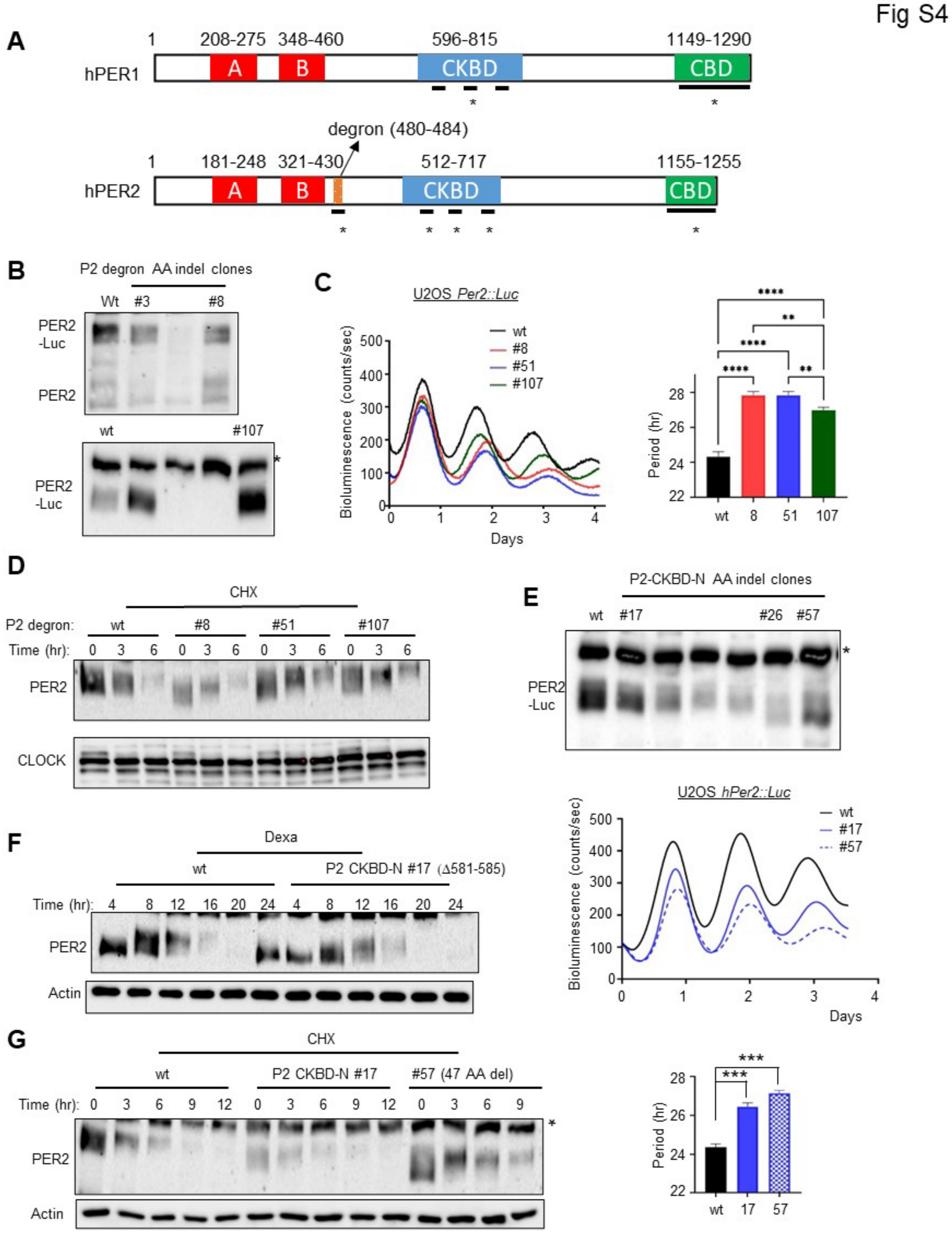

**Fig S4. Gross phosphorylation is not affected when disruptions are made in degron motif or CKBD-N domain, related to Fig 4.**
(A) A schematic diagram of disruptions made in PER domains. Asterisks indicate AA indels generated by CRISPR.
(B-D) PER2 phosphorylation is not affected by AA indels in the degron motif. Circadian rhythms are lengthened in these mutants consistent with the degron mutant mouse model [3]. n=3, mean+/-SEM, * p<0.05, ** p<0.01, *** p<0.001, ****p<0.0001.
(E-G) AA indels in CKBD-N do not compromise phosphorylation and functionality of PER2. PER2 oscillates and gets hyperphosphorylated as much as wt PER2 in various conditions. n=3, mean+/-SEM, * p<0.05, ** p<0.01, *** p<0.001.

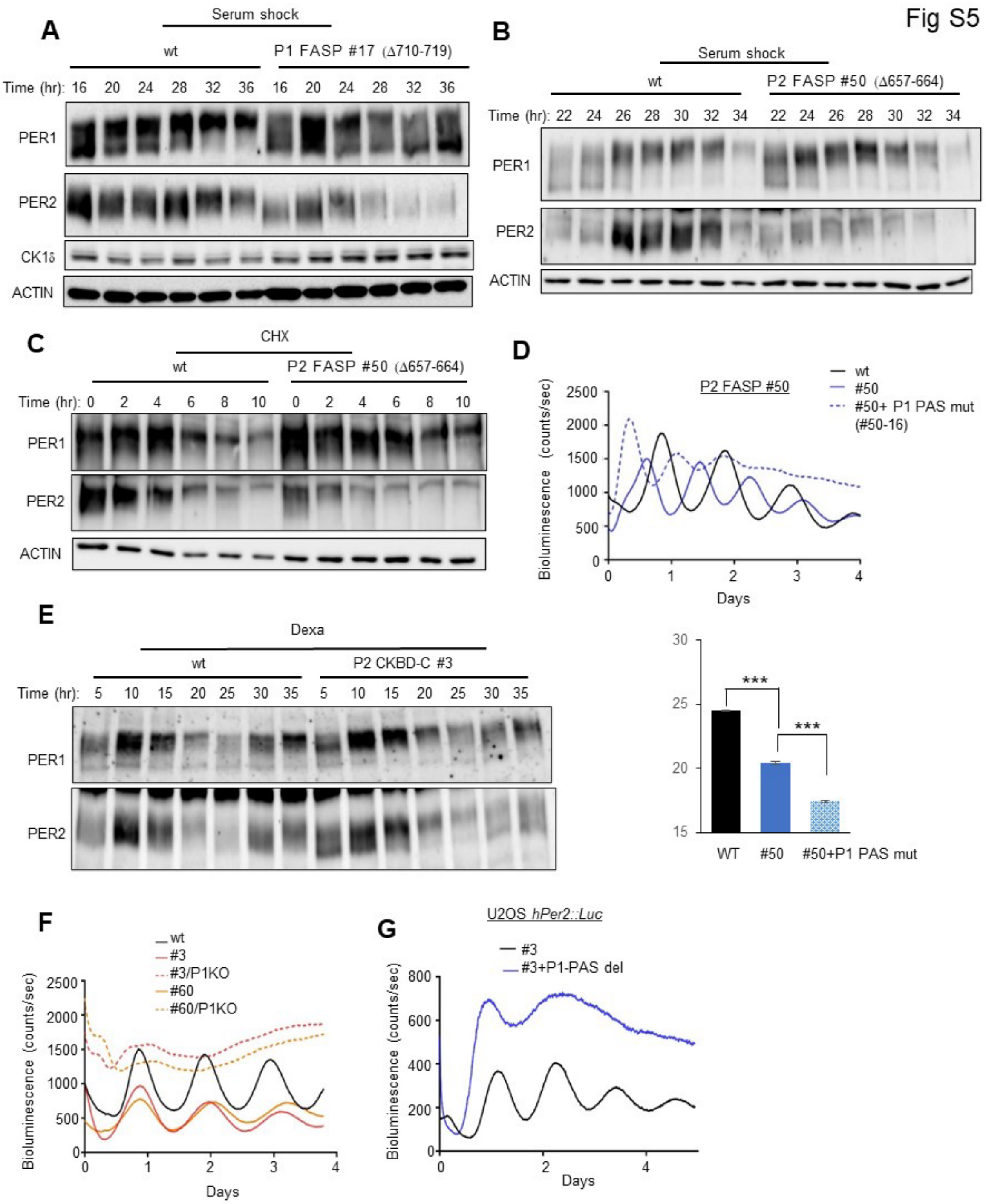

**Fig S5. Gross phosphorylation is not affected by AA indels in the middle of CKBD domain (FASP) while it was slightly reduced when disruptions are made in CKBD-C, related to Fig 4.** (A-C) Immunoblots of protein extracts from mutant U2OS cell lines show that AA indels in the FASP motif do not disrupt gross phosphorylation in PER1 and PER2. (D) A PER2 FASP mutant is functional: circadian bioluminescence rhythms occur, though with shortened period when *Per1* is not functional. n=3, mean+/-SEM, * $p<0.05$, ** $p<0.01$, *** $p<0.001$. (E-G) PER2 CKBD-C mutants can be phosphorylated and oscillate in the presence of wt PER1, suggesting that hetero-dimerization through PER1-PAS enables trans-phosphorylation.

Fig S6

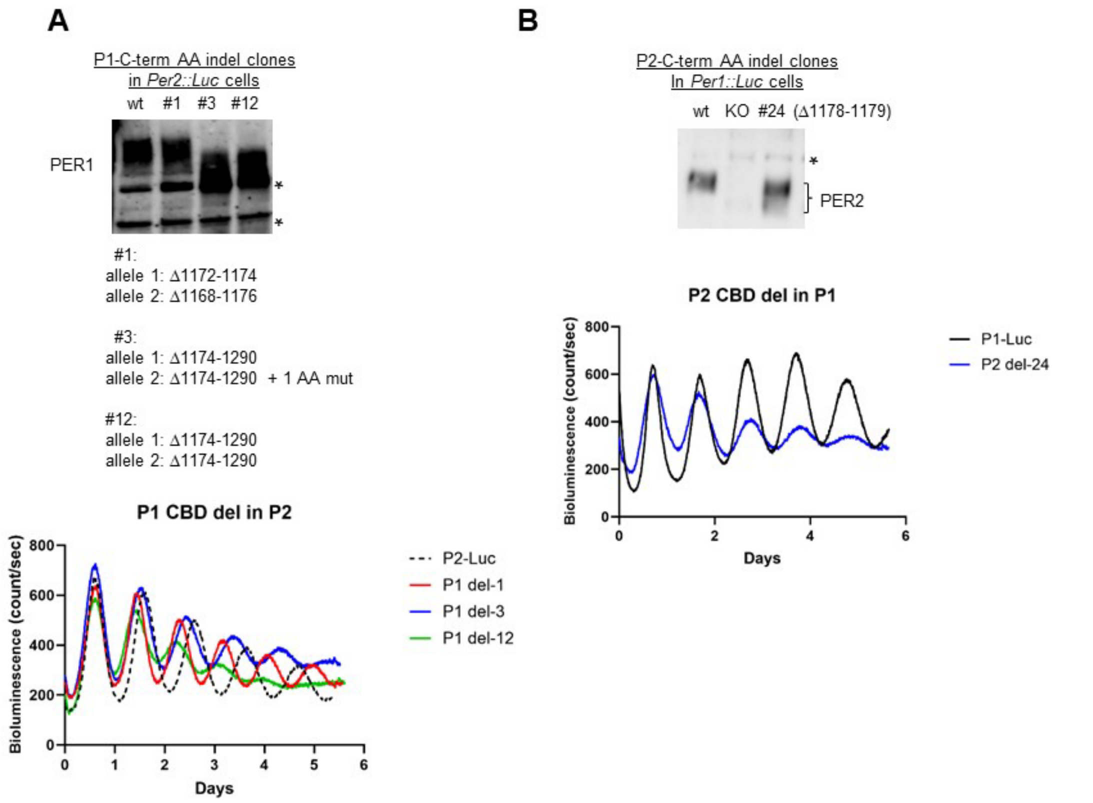

**Fig S6. Gross phosphorylation is not affected by AA indels in CBD, related to Fig 4.** (A and B) Immunoblots of mutant proteins show similar levels of gross phosphorylation as wt PER. Both in-frame and frame-shift mutants are shown. Circadian function in reporter cells is also largely retained. PER1 and PER2 CBD were targeted by sgRNA: gCGCTGGTCCTCAGAAAACCG (AGG) and gTTCTGACTCTCCGTGAACCT (GGG), respectively. Representative of three samples each.

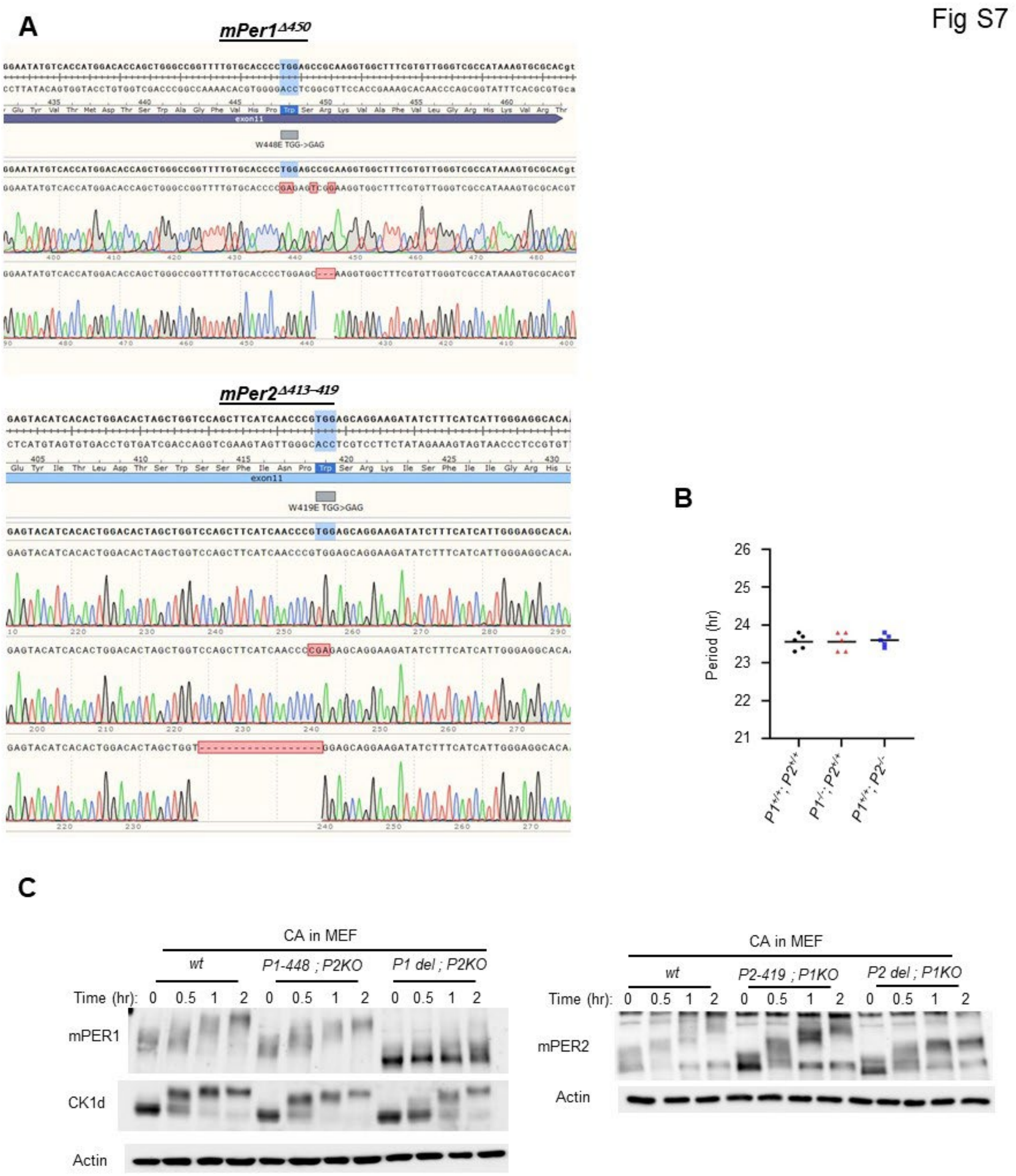

**Fig S7. mPER PAS-B deletion mutants are inherently defective in phosphorylation like the hPER mutant proteins in U2OS cells, related to Fig. 5.** (A) Sanger sequencing results for *mPer1* and *mPer2* deletion mutants. (B) *mPer1* and *mPer2* are redundant for behavioral rhythms. (C) mPER deletion mutants are not hyperphosphorylated by CA treatment in MEFs. Consistent with the data in U2OS cells, mPER2 deletion mutant is more phosphorylated than mPER1 deletion mutant by CA treatment.

Fig S8

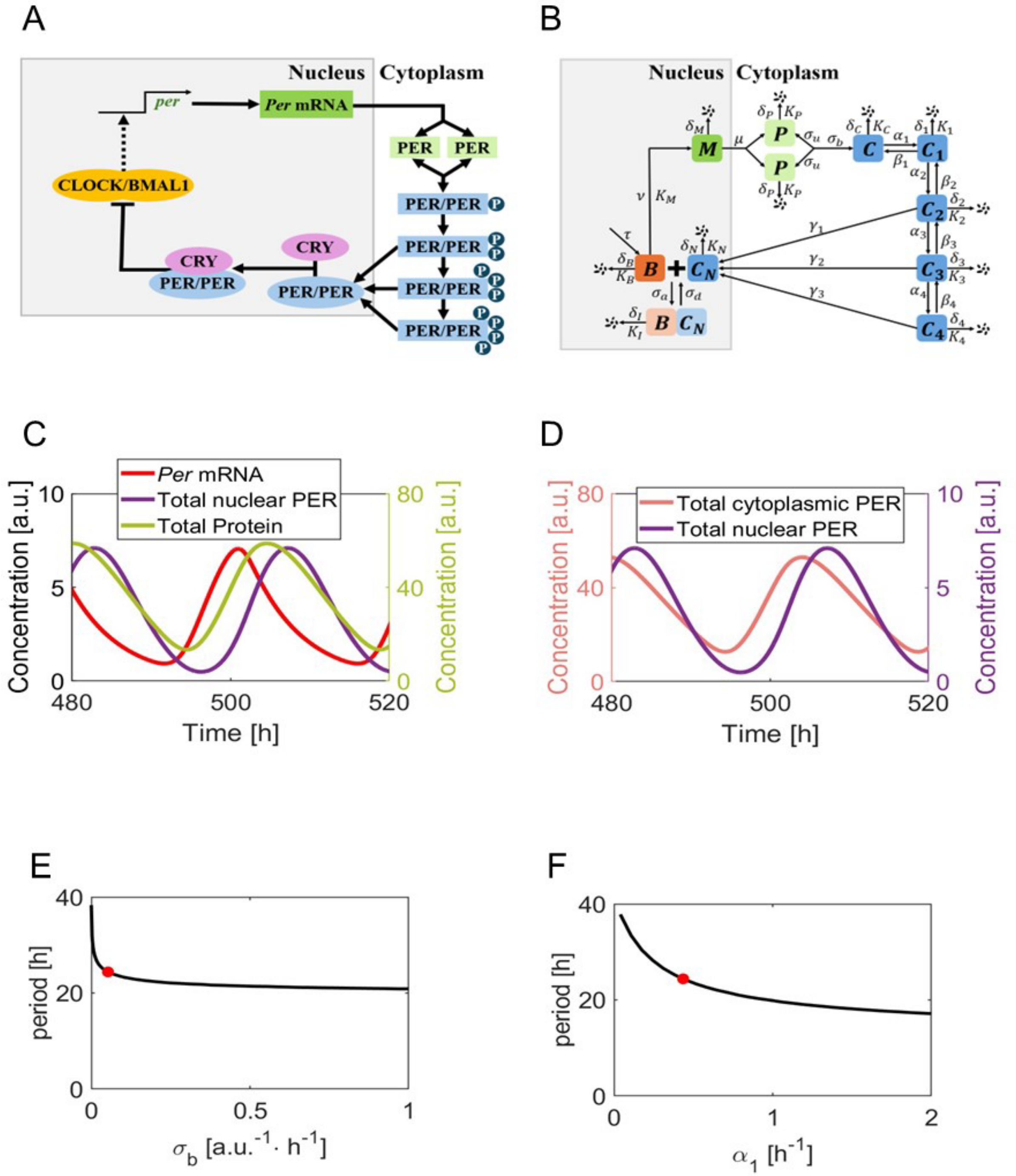

**Fig S8. Wiring diagrams of our mathematical model, related to Fig 7.** (A and B) A diagram of a negative feedback loop with major clock components. CLOCK:BMAL1 ($B$) activates transcription of *Per* mRNA ($M$), which is translated to monomer PER ($P$). Dimerized PER ($C$) would then undergo progressive phosphorylation ($C_1, \cdots, C_4$), and the multi-phosphorylated PER proteins translocate into the nucleus ($C_N$). Upon entering the nucleus, PER would bind to CLOCK:BMAL1 which makes an inactive complex ($I$), inhibiting its own transcription. (C-D) Time evolution of variables in the model with the default values of parameters (Table 1). Here, Total nuclear PER is the sum of nuclear PER that are bound ($I$) or not bound ($C_N$) to CLOCK:BMAL1, total protein is the sum of all the PER proteins ($P, C, C_1, \cdots, C_4, C_N$, and $I$), and total cytoplasmic PER is the sum of all cytoplasmic PER species ($P, C, C_1, \cdots, C_4$). (E-F) Periodograms of parameter $\sigma_b$ and $\alpha_1$. Solid black curves represent the period of oscillations as we vary each parameter. The default value of each parameter is marked with red filled circle, where the period is approximately 24 hrs.

### Table S1. Cell lines used in this study

| Cell lines | Target | gRNA Sequences (PAMs) | **Mutant clones**<br>**Underlined: KO (out of frame)**<br>**Bold: sequenced, in-frame alleles are shown** |
|---|---|---|---|
| U2OS Bm1-Luc | hPer1 PAS A | TGCAAGCGGGACGTGTTCCG(GGG) | 1-3, 2-4, ***2-12***, 2-13 |
| 231 260<br>wt IVYISEQAAVLLRCKRDVFRGTRFSELLAP<br>2-12 IVYISEQAAVLLR---------RFSELLAP (Δ244-252) |  |  |  |
| U2OS Bm1-Luc/Per2 KO #12 | hPer1 PAS A | TGCAAGCGGGACGTGTTCCG(GGG) | ***5***, ***19***, **20**, 40, 44 |
| 221 270<br>wt SVAVSFLTGRIVYISEQAAVLLRCKRDVFRGTRFSELLAPQDVGVFYGST<br>5 SVAVSFLTGRIVYISEQAA-----------GTRFSELLAPQDVGVFYGST (Δ240-250)<br>20 SVAVSFLTGRI-----------------------------------YGST (Δ232-266) |  |  |  |
| U2OS Bm1-Luc | hPer1 PAS B | AAGGCTACCTTGCGGCTCCA(GGG) | **7**, 32, 43, 56, ***72*** |
| 441 480<br>wt WAGFVHPWSRKVAFVLGRHKVRTAPLNEDVFTPPAPSPAP<br>7 WAGFVHP-----------------------------SPAP (Δ448-476)<br>72 WAG------RKVAFVLGRHKVRTAPLNEDVFTPPAPSPAP (Δ443-449) |  |  |  |
| U2OS Bm1-Luc/Per2 KO #12 | hPer1 PAS B | AAGGCTACCTTGCGGCTCCA(GGG) | ***16***, 21 |
| 441 460<br>wt WAGFVHPWSRKVAFVLGRHK<br>16 WAGFV----------LGRHK (Δ446-455) |  |  |  |
| U2OS hPer1::Luc | hPer1 PAS B | AAGGCTACCTTGCGGCTCCA(GGG) | ***11***, 21, 13,21, ***22***, ***23***, ***32***, ***33*** |
| 421 460<br>wt DHSPIRFCARNGEYVTMDTSWAGFVHPWSRKVAFVLGRHK<br>11 DHSPIRFCA----------------------------RHK (Δ430-457)<br>22 DHSPIRFCARNGEYVTMDTSWAGFVH ------FVLGRHK (Δ447-453)<br>23 DHSPIRFCARNGEYVTMDTSWAGFVHPW-----***I***VLGRHK (Δ449-453)<br>32 DHSPIRFCARNGEYVTMDTSWAGFVHPW---***I***AFVLGRHK (Δ449-451)<br>33 DHSPIRFCARNGEYVTMDTSWAG------RKVAFVLGRHK (Δ444-449) |  |  |  |
| U2OS hPer1::Luc | hPer1 FASP | GCCAATAAGGCGGAGAGTG (TGG) | ***14***, *17*, *38* |

<table>
<tr><td colspan="4">671<br>wt GPVSVGTKKDPPSAALSGEGATPRKEPVVGGTLSPLALANKAESVVSVTSQCSFSSTIVH<br>14 GPVSVGTKKDPP------------------------------------------------<br><br>wt VGDKKPPESDIIMMEDLPGLAPGPAPSPAPSPTVAPDPAPDAYRPVGLTKAVLSLHTQKE<br>14 ------------------------------------------------------------<br><br>wt EQAFLSRFRDLGRLRGLDSSSTAPSALGERGCHHGPAPPSRRHHCRSKAKRSRHHQNPRA<br>14 ------------------------------------------------------------<br>890<br>wt EAPCYVSHPSPVPPSTPWPTPPATTPFPAVVQPYPLPVFS<br>14 ----------------------------AVVQPYPLPVFS (Δ683-878)</td></tr>
<tr><td>U2OS Bm1-Luc<br>(P1KO)#3</td><td>hPer2 PAS A</td><td>ATCTCTTTTACAGTGAAATA(TGG)</td><td>28, 112</td></tr>
<tr><td colspan="4">201 230<br>wt SGKILYISDQVASIFHCKRDAFSDAKFVEF<br>28 SGKI---------------------KFVEF (Δ205-225)<br>112_1 SGKILYISD-----------AFSDAKFVEF (Δ210-220)<br>112_2 SGKILYISD------HCKRDAFSDAKFVEF (Δ210-215)</td></tr>
<tr><td>U2OS hPer1::Luc</td><td>hPer2 PAS A</td><td>ATCTCTTTTACAGTGAAATA(TGG)</td><td><u>7</u>, 12, <u>55</u></td></tr>
<tr><td colspan="4">203 249<br>wt KILYISDQVASIFHCKRDAFSDAKFVEFLAPHDVGVFHSFTSPYKLP<br>12 KILYISD-------------------------------------<u>Q</u>LP (Δ210-246)</td></tr>
<tr><td>U2OS hPer2::Luc</td><td>hPer2 PAS B</td><td>GCTTCATCAACCCATGGAGC(AGG)</td><td>2, 5, 8, 13, 36, 52, 53, 55, 58</td></tr>
<tr><td colspan="4">411 430<br>wt DTSWSSFINPWSRKISFIIG<br>2 DTSWSSFI<u>T</u>-WSRKISFIIG (Δ420)<br>5 DTSWSSF<u>T</u>--WSRKISFIIG (Δ419-420)<br>8 DTSWSSFIN---RKISFIIG (Δ420-422)<br>13 DTSWSSFI<u>K</u>-WSRKISFIIG (Δ420)<br>36 DTSWSSFI<u>R</u>----KISFIIG (Δ420-423)<br>52 DTSWSSFI<u>R</u>----KISFIIG (Δ420-423)<br>53 D--------------SFIIG (Δ412-425)</td></tr>
<tr><td>hPer2 PAS B mutant<br>#52/Per1 KO</td><td>hPer1 PAS B</td><td>AAGGCTACCTTGCGGCTCCA(GGG)</td><td>52-100</td></tr>
<tr><td colspan="4">411 430<br>wt DTSWSSFINPWSRKISFIIG</td></tr>
</table>

<table>
<tr><td colspan="4">52-100 DTSWSSFI<u>R</u>----KISFIIG (Δ420-423) + Per1 KO (frame-shift in hPer1 PAS B)</td></tr>
<tr><td>U2OS hPer2::Luc</td><td>FASP</td><td>GCCGGGCAAGGCAGAGAGTG(TGG)</td><td>50</td></tr>
<tr><td colspan="4">651 670<br>wt LTSLALPGKAESVASLTSQC<br>50_1 LTSLALPGK------LTSQC (Δ660-665)<br>50_2 LTSLALP--------LTSQC (Δ658-665)</td></tr>
<tr><td>hPer2 FASP #50/Per1 PAS A mutant</td><td>hPer1 PAS A</td><td>TGCAAGCGGGACGTGTTCCG(GGG)</td><td>50-16</td></tr>
<tr><td colspan="4">220 270<br>Wt (hPER1) LTSLSVAVSFLTGRIVYISEQAAVLLRCKRDVFRGTRFSELLAPQDVGVFYGSTALPGKL<br>50-16 LTSLSV----------------------------------------GVFYGSTALPGKL</td></tr>
<tr><td>U2OS hPer2::Luc</td><td>Degron</td><td>CACAGCGGCTCCAGTGGCTA(CGG)</td><td>3, 8, 14, 51, 52, 107</td></tr>
<tr><td colspan="4">471 490<br>wt LQPVPHSGSSGYGSLGSNGS<br>3 LQPVPH-----------NGS (Δ477-487)<br>8 LQPVPHS----<u>D</u>GSLGSNGS (Δ478-481)<br>51 LQPVPHSGS--YGSLGSNGS (Δ480-481)<br>107 LQPVPHSG---YGSLGSNGS (Δ479-481)</td></tr>
<tr><td>U2OS hPer2::Luc</td><td>CKBD-N</td><td>TGTCCAAGCAGCTGATCTGC (TGG)</td><td>17, 26, 33, 57</td></tr>
<tr><td colspan="4">571 590<br>wt LACKNQPTCSYQQISCLDSV<br>17 LACKNQPTCS-----CLDSV (Δ581-585)<br>26 Not sequenced<br>57 LACKNQPTCSYQQ---------- (Δ584-633+3AA=47 AA del)</td></tr>
<tr><td>U2OS hPer2::Luc</td><td>CKBD-C</td><td>CAGCGAGTACCTCCTTGGTG (AGG)</td><td>3, 38 60, 46, 127-3</td></tr>
<tr><td colspan="4">711 740<br>wt CGLSQEKEPFKKLGLTKEVLAAHTQKEEQS<br>3 CGLSQEKEPFK------------------<u>N</u> (Δ722-739)<br>60 CGLSQEKEPFKKL-----VLAAHTQKEEQS (Δ724-728)<br>127(3) CGLSQEKE-------------------EQS (Δ719-737)</td></tr>
<tr><td>U2OS hPer2::Luc<br><br>Per1 KO in #3, #60</td><td>hPer1 PAS B</td><td>AAGGCTACCTTGCGGCTCCA(GGG)<br><br>Per1KO confirmed by WB</td><td>3-34, 60-17</td></tr>
<tr><td colspan="4">711 740<br>wt CGLSQEKEPFKKLGLTKEVLAAHTQKEEQS<br>3-34 CGLSQEKEPFK------------------<u>N</u> (Δ722-739) + P1KO<br>60-17 CGLSQEKEPFKKL-----VLAAHTQKEEQS (Δ724-728) + P1KO</td></tr>
</table>

<table>
<tr><td>U2OS<br>*hPer2::Luc*<br><br>P1 PAS-B<br>mutant in #3</td><td>hPer1 PAS B</td><td>AAGGCTACCTTGCGGCTCCA(GGG)<br><br>PAS-B del confirmed by WB</td><td>***3-15***</td></tr>
<tr><td colspan="4">     711                          740<br>wt     CGLSQEKEPFKKLGLTKEVLAAHTQKEEQS<br>3-15   CGLSQEKEPFK------------------***N*** (Δ722-739) + P1 PAS-B del</td></tr>
</table>
